# Genomics of an avian neo-sex chromosome reveals the evolutionary dynamics of recombination suppression and sex-linked genes

**DOI:** 10.1101/2020.09.25.314088

**Authors:** Hanna Sigeman, Maria Strandh, Estelle Proux-Wéra, Verena E. Kutschera, Suvi Ponnikas, Hongkai Zhang, Max Lundberg, Lucile Soler, Ignas Bunikis, Maja Tarka, Dennis Hasselquist, Björn Nystedt, Helena Westerdahl, Bengt Hansson

## Abstract

How the avian sex chromosomes first evolved from autosomes remains elusive as 100 million years (Myr) of divergence and degeneration obscure their evolutionary history. Sylvioidea songbirds is an emerging model for understanding avian sex chromosome evolution because a unique chromosome fusion event ∼24 Myr ago has formed enlarged “neo-sex chromosomes” consisting of an added (new) and an ancestral (old) part. Here, we report the female genome (ZW) of one Sylvioidea species, the great reed warbler (*Acrocephalus arundinaceus*). We confirm that the added region has been translocated to both Z and W, and show that the added-W part has been heavily reorganised within itself and with the ancestral-W. Next, we show that recombination between Z and W continued after the fusion event, and that recombination suppression took ∼10 Myr to be completed and arose locally and non-linearly along the sex chromosomes. This pattern is inconsistent with that of large inversions and instead suggests gradual and mosaic recombination suppression. We find that the added-W mirrors the ancestral-W in terms of repeat accumulation, loss of genetic variation, and gene degeneration. Lastly, we show that genes being maintained on W are slowly evolving and dosage sensitive, and that highly conserved genes across broad taxonomic groups, regardless of sex-linkage, evolve slower on both Z and W. This study reveals complex expansion of recombination suppression along avian sex chromosomes, and that the evolutionary trajectory of sex-linked genes is highly predictable and governed partly by sex-linkage *per se*, partly by their functional properties.

## 1. INTRODUCTION

Sex chromosomes have evolved from autosomes many times across the animal and plant kingdoms and have been studied intensely for their role in sex determination but also for their other unique characteristics, such as loss of recombination and sex-specific evolutionary pressures (Bachtrog et al. 2011; Abbott et al. 2017). Traditionally, most research on sex chromosomes has been done on species with highly heteromorphic sex chromosomes. However, the degeneration of such old sex chromosomes obscures the genomic signatures of their early evolutionary history. To learn more about the transition of sex chromosomes from their autosomal origin, newly formed sex chromosomes (formed *de novo* or by turnovers) or partially newly formed sex chromosomes (neo-sex chromosomes) are more suitable systems (Wright et al. 2016; Ponnikas et al. 2018).

In birds, the sex chromosomes (Z and W) originated more than 100 Myr ago (Zhou et al. 2014) as recombination became suppressed around the sex-determining gene (*DMRT1*; Smith et al. 2009). Since then, the sex chromosome copies have ceased to recombine along most of their length in the majority of species, except in some paleognaths (e.g. ratites and tinamous), resulting in heavy differentiation between Z and W with weak signatures of their shared origin and few surviving genes on the degenerated sex-limited W chromosome (Zhou et al. 2014; Smeds et al. 2015; Bellott et al. 2017). Birds have highly stable karyotypes with few inter-chromosomal rearrangements compared to other vertebrates (Ellegren 2010), and the Z chromosome has been shown to share synteny across its entire length even between widely different clades of birds (Nanda et al. 2008). However, there are some exceptions, all so far found among passerines, where autosome–sex chromosome fusions have enlarged the original sex chromosomes and formed neo-sex chromosomes (Pala et al. 2012; Sigeman et al. 2019; Gan et al. 2019; Dierickx et al. 2020; Sigeman et al. 2020). Such events often lead to an extension of recombination suppression to include also the translocated chromosomal region, which then becomes bound by the same evolutionary processes as the original sex chromosome (Bachtrog 2013). These neo-sex chromosomes provide excellent opportunities to study the drivers of recombination suppression between avian sex chromosomes and allow us to study rates of evolution between sex-linked genetic regions of different ages.

The songbird superfamily Sylvioidea (*sensu lato*; Moyle et al. 2016; Oliveros et al. 2019) split from other songbirds around 24 Myr ago and has since undergone one of the fastest radiations within birds, with over 1,200 extant species (Alström et al. 2006). All Sylvioidea birds studied so far, i.e. species representatives of 11 of the 22 families within Sylvioidea (Pala et al. 2012; Sigeman et al. 2019; Sigeman et al. 2020; Leroy et al. 2019), share a unique karyotype feature: a neo-sex chromosome formed by a chromosomal fusion between the ancestral sex chromosomes and a part of chromosome 4A (according to chromosome naming from the zebra finch, *Taeniopygia guttata;* Warren et al. 2010). This fusion has thus added new genomic material to the sex chromosomes of Sylvioidea birds, characterized by less Z-to-W differentiation and W degeneration compared to the ancestral part. Here, we present a detailed study of the evolutionary history of this neo-sex chromosome in a Sylvioidea species belonging to the family Acrocephalidae, the great reed warbler (*Acrocephalus arundinaceus*). By constructing a high-quality annotated reference genome from a female great reed warbler, containing both a Z and W chromosome, we are able to study both the chromosomal structure of this fusion event and how the previously autosomal region has evolved in this novel sex-linked environment in terms of recombination suppression, repeat accumulation and gene differentiation.

## 2. RESULTS

### 2.1 Sequencing, assembly, annotation and synteny

We sequenced high molecular weight DNA of a female great reed warbler from our long-term study population in southern central Sweden using a combination of long-read, linked-read, short-read sequencing, and optical data, to reconstruct its genome *de novo* (see details on raw data in Supplementary Table 1a and genome statistics correlating to each stage in the genome assembly process in Supplementary Tables 2 and 3). We also used information from a linkage map analysis (Ponnikas et al. 2020), based on a multi-generation pedigree of great reed warblers genotyped with Restriction site–Associated DNA (RAD) sequencing (Hansson et al. 2018), to identify and correct assembly errors (see Methods). The final assembly (acrAru1) consisted of 3,013 scaffolds and had an N50 of 21.4 Mb (Figure 1; Supplementary Table 2). The number of conserved avian single-copy orthologs (*n* = 4,915) was assessed with BUSCO v.3.0.2 (aves_odb9 dataset; Simão et al. 2015). The final assembly had 93.1% complete genes (Supplementary Table 3), which is similar to other long-read sequenced genomes (e.g. between 93% and 94% in five species of birds-of-paradise, family Paradisaeidae; Xu et al. 2019). The total repeat content of the final draft assembly was 19%, with LTR elements as the most common type of repeat (7.9%) followed by LINEs (5.5%; Supplementary Table 4). The genome assembly was annotated with 22,524 genes. Scaffolds belonging to the non-recombining part of the Z and the W chromosome, respectively, were detected by evaluating sex-specific differences in read coverage and/or heterozygosity (Supplementary Table 5) using whole-genome resequencing data from five male (ZZ) and five female (ZW) great reed warblers (Supplementary Table 1b) which were aligned to the genome assembly.

**Figure 1.**
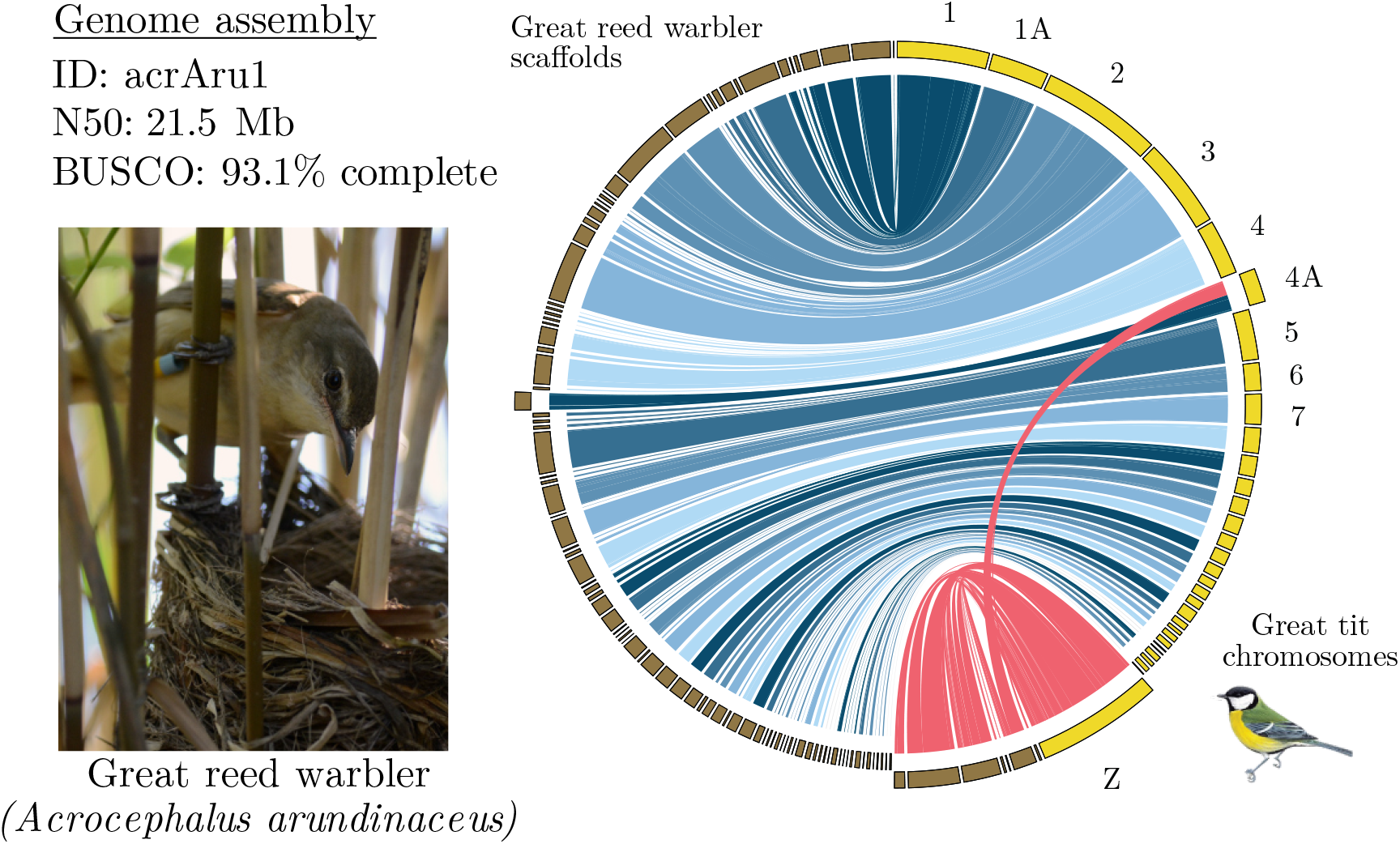
The great reed warbler and its genome assembly. Synteny analysis between the great reed warbler (scaffolds in brown) and the great tit genome (chromosomes in yellow) showing a single inter-chromosomal rearrangement: the autosome–sex chromosome fusion unique to Sylvioidea songbirds involving the Z chromosome and a 9.6 Mb part of chromosome 4A. Links involving sex-linked great reed warbler scaffolds are red, and links involving autosomal scaffolds are coloured in different shades of blue. Great reed warbler scaffolds that share synteny (filtered to exclude short matches, see Methods) with great tit chromosomes Z and 4A, and these two chromosomes, are scaled to twice their actual size for illustrative purposes. Photograph of a female great reed warbler by August Thomasson.

A comparison between the genomes of the great reed warbler and the great tit (*Parus major*), which is the closest relative to the great reed warbler with a near-complete chromosome-level assembly (lacking only a few microchromosomes and the W chromosome), showed largely conserved synteny (Figure 1). We detected a single well-supported inter-chromosomal rearrangement: the autosome–sex chromosome fusion unique to Sylvioidea songbirds involving chromosome Z and approximately half (0-9.6 Mb) of chromosome 4A (20.7 Mb in total; Figure 1) through a scaffold bridging over between these two chromosome regions (see also below). This confirms the fusion between chromosome Z and chromosome 4A that has occurred basally in the Sylvioidea clade (Pala et al. 2012; Sigeman et al. 2020). The intra-chromosomal collinearity between the species was disrupted for several macro- as well as microchromosomes (Supplementary Figure 1). Throughout this paper we will refer to the great reed warbler neo-sex chromosome region sharing synteny with other songbird Z chromosomes as the ancestral sex chromosome region (abbreviated as ancestral-Z and -W for the two sex chromosome copies, respectively), and the translocated region sharing synteny with chromosome 4A as the added sex chromosome region (abbreviated as added-Z and -W, respectively).

### 2.2 Sex chromosome structure and cross-species homology

We identified 22 Z-linked scaffolds (total length of 88.7 Mb; mean length 4 Mb; Supplementary Table 5) in the great reed warbler genome. A detailed comparison to the zebra finch genome showed that these scaffolds share synteny with either the ancestral Z chromosome or the first part of chromosome 4A, with the exception of one scaffold (Scaffold31; Figure 2a; Supplementary Table 6) that shares synteny with the end of chromosome Z (position 67.6-72.9 Mb) and a large part of chromosome 4A (position 9.6-0.9 Mb). This locates the fusion point in the zebra finch genome to chromosome Z position 72.9 Mb and chromosome 4A position 9.6 Mb (Figure 2c; Pala et al. 2012; Sigeman et al. 2020).

**Figure 2.**
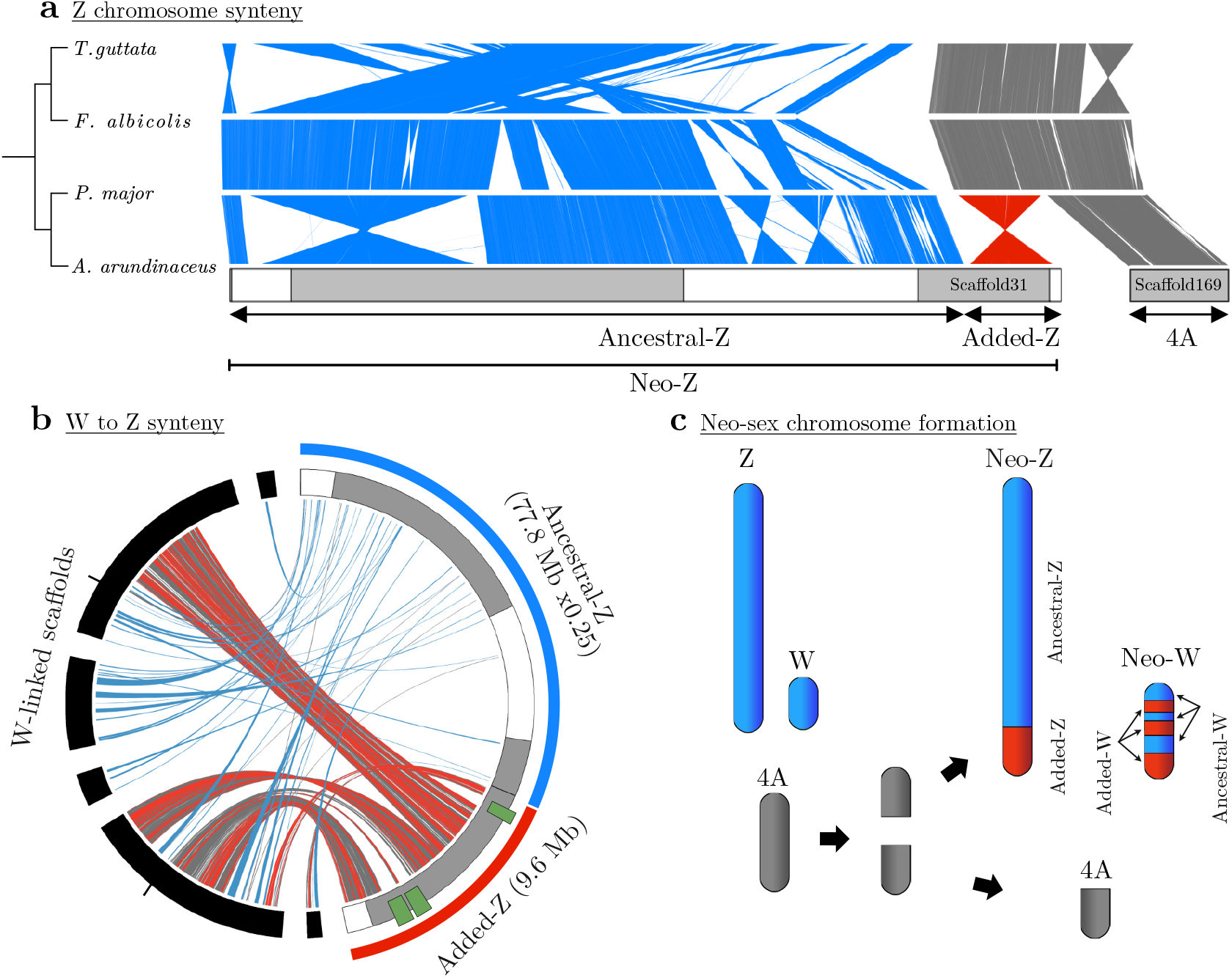
Structure of the great reed warbler Z and W neo-sex chromosomes. **(a)** Chromosome alignments of chromosome Z (blue) and 4A (grey) in four species of passerines, with the part of 4A representing the added-Z region in the great reed warbler indicated (red). Phylogenetic relationships between the species are depicted in a cladogram (left), great reed warbler scaffold lengths are outlined and Scaffold31, crossing the fusion point between the ancestral-Z and added-Z chromosome regions, is indicated. Scaffold169, which aligns with chromosome 4A and segregates as a separate autosomal chromosome, is also indicated. **(b)** Syntenic regions between the great reed warbler W-linked (black) and Z-linked (grey and white alternating) scaffolds. The Z-linked scaffolds belonging to the ancestral sex chromosome region (blue) are scaled to 25% of their true size for illustrative purposes. Two pairs of W scaffolds shown to be physically linked (by linked-read data from a different female; see Main text) are placed next to each other, without gaps but with tick marks at the scaffold boundaries. The grey links show syntenic information on a genomic level, while chromosomal positions of gametologous (ZW) gene pairs are shown as blue links (for ancestral sex chromosome genes) and red links (added sex chromosome genes). Note that four W-linked scaffolds have genes with orthologs on both the ancestral and added sex chromosome regions, strongly suggesting that the ancestral-W and added-W are physically connected. Green symbols mark putative W-deletions (see Main text). **(c)** A graphic representation of the fusion event forming the Sylvioidea neo-sex chromosome system.

Moreover, six of the 22 Z-linked scaffolds, which together cover 98.6% of the total length of the Z-linked scaffolds, were possible to order with our RAD-based linkage map analysis (Figure 2a; Supplementary Table 7; also see Ponnikas et al. 2020). Linkage mapping further identified a scaffold (Scaffold217; 0.9 Mb in length) containing the pseudoautosomal region (PAR), i.e. the region where the Z and W chromosomes recombine (see Ponnikas et al. 2020). Alignments to the genomes of zebra finch, collared flycatcher (*Ficedula albicollis*) and great tit confirmed that the great reed warbler Z chromosome consists of an initial part corresponding to the PAR (0-0.9 Mb; which is not included in the Z chromosome sequence of the other species and thus not shown here), a large central part corresponding to the ancestral Z (0.9-77.8 Mb), and a final part, the added region, corresponding to the first half (9.6 Mb) of chromosome 4A (77.8-87.5 Mb; Figure 2a; Supplementary Table 6,7). The linkage map analysis further showed that the scaffold corresponding to the second half of chromosome 4A (Scaffold169; 9.6-20.7 Mb) segregates autosomally in the great reed warbler (Figure 2a), i.e. confirming the fission of chromosome 4A in Sylvioidea (Figure 2c). In addition to the fusion between the ancestral Z and chromosome 4A in Sylvioidea, three large inversions broke the collinearity between the great reed warbler and the great tit and the collared flycatcher Z chromosomes, whereas the zebra finch differed by several Z chromosome rearrangements as previously described (Itoh et al. 2006; Kawakami et al. 2014) (Figure 2a).

The 50 W-linked scaffolds had a total length of 30.2 Mb (mean length 0.6 Mb). Of these, 15 scaffolds (with a total length of 28.0 Mb; mean length 1.9 Mb) were present in the gene annotation while the remaining ones were short and not present (Supplementary Table 5). We identified and manually curated 147 gametologous (ZW) gene pairs in the great reed warbler gene annotation; 41 pairs from the ancestral and 106 from the added sex chromosome region (Supplementary Table 8). Of the 41 ancestral W genes, 36 had previously been described in a detailed study of the W chromosome in the collared flycatcher (Smeds et al. 2015; Supplementary Table 9). The collared flycatcher annotation included an additional eight W genes of which six were present in our great reed warbler annotation under the same gene name as in the flycatcher but did not have enough evidence to be classified as orthologs and were therefore not included here (see Methods). We aligned the 15 W-linked scaffolds to the great reed warbler Z-linked scaffolds, and cross-positioned the 147 ZW gametologs, and found that four great reed warbler W scaffolds showed substantial sequence similarity with, and contained many genes with gametologs on both the ancestral-Z and added-Z chromosome regions, whereas the remaining W scaffolds only contained sequences with similarity to ancestral-Z (Figure 2b). The shared homology of four W-linked scaffolds to both ancestral-Z and added-Z strongly supports that the added-W region has fused with the ancestral-W region, and has subsequently been intra-chromosomally rearranged (Figure 2b,c). Evidence from a *de novo* assembly based on linked-read data from another great reed warbler female (Supplementary Table 1b,2,3) provided independent evidence for the correctness of three of the four scaffolds sharing synteny with both the ancestral-Z and added-Z chromosome regions. This confirms the presence of a single enlarged W chromosome in Sylvioidea birds (consisting of the ancestral W plus a part of chromosome 4A) as opposed to the alternative of two separate W chromosomes (i.e. a ZW_1_W_2_ system). The synteny analyses between the great reed warbler Z and W further showed three larger deletions on the W chromosome with a total size of ∼2.4 Mb (Figure 2b). Despite these deletions, the total length of great reed warbler W scaffolds with synteny to added-Z (10.6 Mb) is larger than the corresponding Z scaffolds (9.6 Mb). To conclude, our data clearly support a fusion of the ancestral songbird sex chromosome and a part of chromosome 4A (inverted) in the great reed warbler, and that this fusion involves both Z and W, forming young (∼24 Myr; see below) and enlarged Z and W neo-sex chromosomes.

### 2.3 Evolution of recombination suppression

Our genomic and linkage map data show that the neo-sex chromosomes have ceased to recombine over most of the ancestral part (except the PAR) and over the whole added region in present-day great reed warblers. To estimate when the different parts of the great reed warbler Z and W chromosomes ceased to recombine, we used a phylogenetic approach to place each great reed warbler W gametolog copy on gene trees with a fixed and dated topology (see Methods). The phylogeny included five Sylvioidea species being short-read sequenced in this study (Supplementary Table 1b), and six non-Sylvioidea birds and the green anole (*Anolis carolinensis*) with available sequence data, selected to widely represent divergence times to the great reed warbler. We dated the nodes in the phylogeny with a set of autosomal gene sequences using four calibration points (see Methods), and estimated the split between Sylvioidea and non-Sylvioidea to ∼24 Myr (Supplementary Figure 2).

For each of the 147 W gametologs on the great reed warbler W chromosome, we searched for orthologous sequences in these species, being either Z-linked (for all genes in the Sylvioidea species and for the ancestral region in the non-Sylvioidea birds) or autosomal (for the added region in the non-Sylvioidea birds and for all genes in the green anole). Sequences from all species were aligned, which resulted in alignments for 24 and 49 genes in the ancestral and added region, respectively. Next, for each gene, we used the evolutionary placement algorithm-approach (EPA) to estimate likelihood weight ratio (LWR) support (see Methods) for the placement of the great reed warbler W sequence at each relevant phylogenetic position, i.e. positions prior to the Sylvioidea radiation for ancestral-W genes, and positions at and after the formation of the Sylvioidea clade for added-W sequences. The power of this analysis depends on the number of informative sites and thus partly on the alignment length. Accordingly, some great reed warbler W genes had several possible placements, i.e. were inconclusive (similar LWR values for different nodes), whereas other W genes showed strong support for a single or few placements (Figure 3). The LWR support for the most likely phylogenetic position of the W gametolog differed among different genes from very high support (>0.95 for 14 genes) to low support (<0.50 for 12 genes; Figure 4b; Supplementary Table 10).

**Figure 3.**
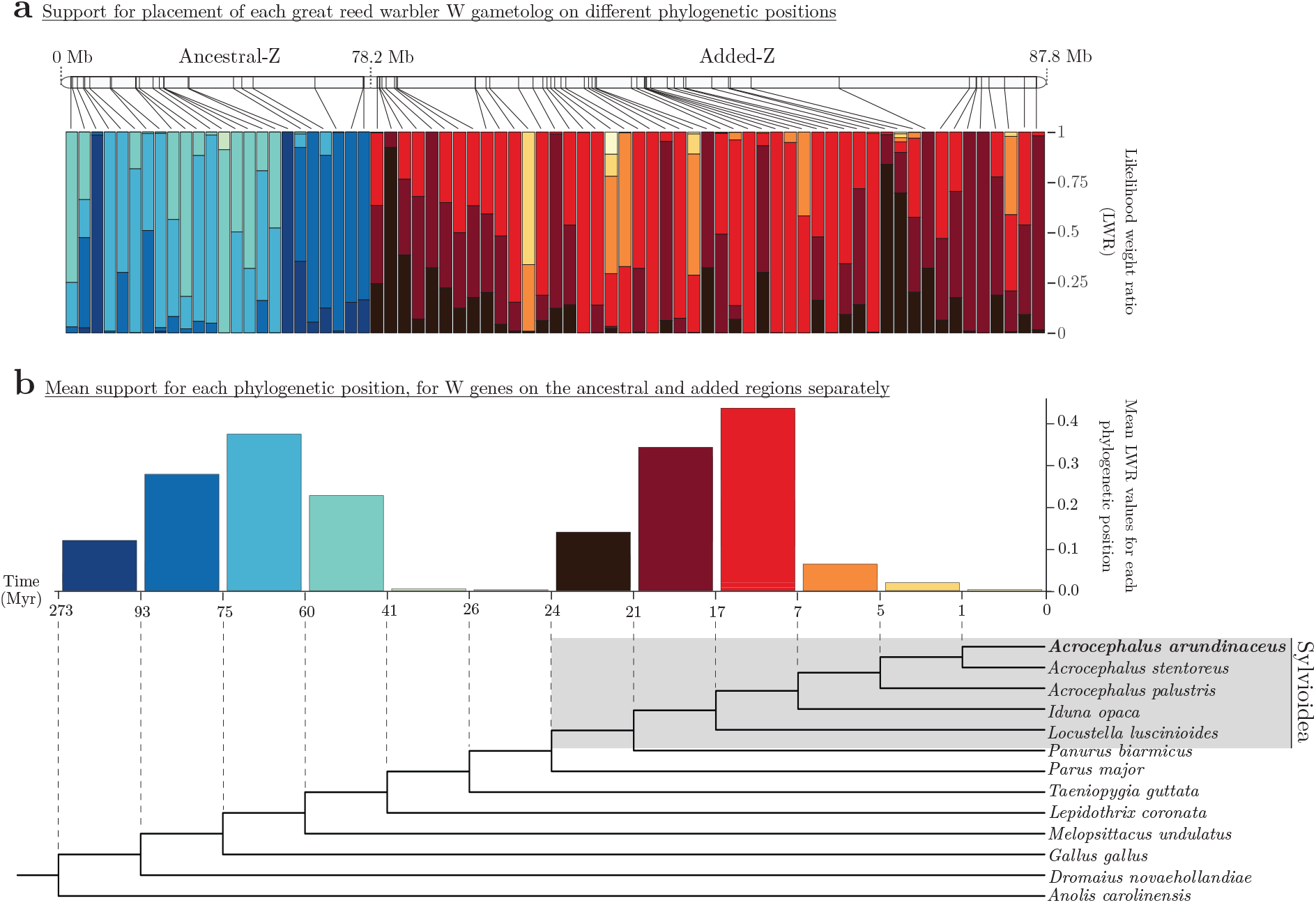
Timing of recombination suppression along the Sylvioidea ancestral-Z and added-Z chromosome regions. **(a)** Stacked bars showing likelihood weight ratio (LWR) values for the placement of each great reed warbler W gametolog on different positions in a fixed and dated avian phylogeny (see (b)) evaluated with evolutionary placement algorithm-analysis. Each stacked bar, corresponding to one gene, shows LWR values for each dated position (one colour per node; see (b)) and adds up to a combined LWR of 1. The chromosome ideogram (above) marks the chromosomal location of each gene on the great reed warbler Z chromosome (the ancestral-Z region with 24 analysed genes, and the added-Z region with 49 analysed genes, are scaled differently for illustrative purposes). **(b)** Bar plot showing mean LWR values for each phylogenetic position for all great reed warbler W gametologs on the ancestral-Z (bars in blue-to-green colours) and added-Z (bars in red-to-yellow colours). The phylogeny shows dated nodes for the great reed warbler and five additional Sylvioidea species, and six non-Sylvioidea birds and the green anole (Anolis carolinensis). We estimated the split between Sylvioidea and non-Sylvioidea to ∼24 Myr (Supplementary Figure 2).

**Figure 4.**
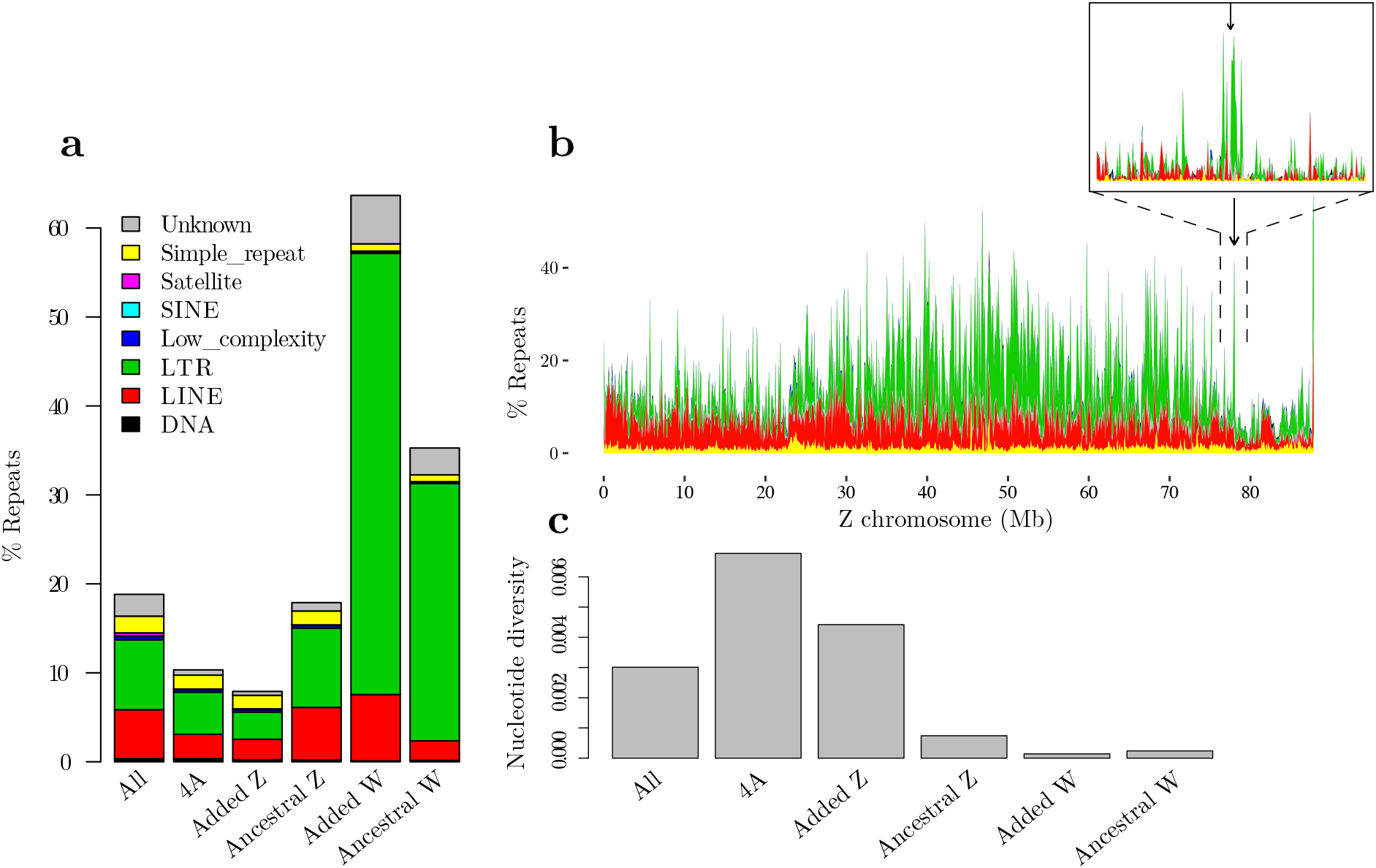
Repeat accumulation and loss of genetic variation on the great reed warbler sex chromosomes. **(a)** Percentages of repeat elements for the entire genome assembly (All), the autosomally segregating part of chromosome 4A (4A), and the added and ancestral parts of Z and W. **(b)** Repeats along the Z chromosome (100 kb windows), following the same colour scheme as Figure 4a. The arrow marks the fusion point between the ancestral and added regions (77.8 Mb). A zoom in of the fusion point with 10 kb window sizes is shown in the top right corner. **(c)** Nucleotide diversity estimates for the genomic regions, calculated from five female great reed warblers.

These results confirm that gametologs on the ancestral sex chromosome ceased to recombine early in the avian phylogeny (cf. Zhou et al. 2014), before the origin of passerines. This was indicated by the placement of all ancestral W gametologs outside passerines, i.e. prior to the split of the suboscine passerine blue-crowned manakin (*Lepidothrix coronata*) ∼41 Myr ago (Figure 3; Supplementary Table 10). Genes with strong support for early (>93-75 Myr ago) and relatively late (60-41 Myr ago) timing of recombination suppression were scattered across the whole ancestral part of the chromosome, with an accumulation of early recombination suppression towards the end half of ancestral-Z (Figure 3). This is in line with previous findings and suggested to be explained by the formation of different layers of recombination suppression, “evolutionary strata”, which since then have become rearranged to different degrees in different lineages (Xu et al. 2019). The timing of recombination suppression that we estimated (ranks were assigned based on the highest LWR value for each gene) for ancestral sex chromosome gametologs correlated significantly with a previously made division into evolutionary strata (strata S0 to S3; Xu et al. 2019) (Spearman correlation: *r*_S_ = -0.75, *n* = 22 genes, *p* = 4.8×10^−5^; Supplementary Table 11). In the much younger added sex chromosome region, genes that ceased to recombine at or early after the fusion event (24-17 Myr ago) were located mainly in the beginning and end of the chromosome region, but also together with genes in the central part that continued to recombine until and even after the formation of Acrocephalidae, represented by *Acrocephalus* and *Iduna*, ∼7 Myr ago (Figure 3). This mosaic and gradual pattern of recombination suppression over several Myr does not support a hypothesis of simple linear progression of recombination suppression along the added sex chromosomes starting from the fusion point.

### 2.4 Repeat accumulation and loss of diversity

Recombination suppression and the following sex-linkage are expected to have severe consequences for the sex chromosomes. In particular the sex-limited copy (Y and W), with its greatly reduced effective population size and lack of recombination in both sexes, is prone to repeat accumulation, loss of genetic variation and gene functionality because of increasing influence of genetic drift and decreasing efficiency of selection (Bachtrog et al. 2011). Our analyses of the great reed warbler genome assembly confirm these patterns. The W scaffolds with synteny to the ancestral sex chromosome region consisted of 64% repeat elements and had the highest proportion of repeats of all chromosomal regions. The corresponding number for W scaffolds with synteny to the added sex chromosome region, with a more recent history of sex-linkage and recombination suppression, was 35% (Figure 4a; Supplementary Table 4). Both of these W regions had a considerably higher repeat content than the genome-wide average (19%), the ancestral-Z scaffolds (18%) and the added-Z scaffolds (8%). The repeat content of the added-Z scaffolds is similar to that of the autosomal part of chromosome 4A (10.4%), suggesting that the repeat landscape of this region has so far been little affected by the translocation to the Z chromosome (Figure 4a). However, the central part of the Z chromosome (25-74 Mb), i.e. the end part of ancestral-Z, which contains the older evolutionary strata previously identified in birds (Zhang et al. 2014; Xu et al. 2019), had a higher proportion of repeat elements than the beginning of the chromosome (0-25 Mb; not including the PAR), and also compared to the added-Z chromosome part with its much more recent history of recombination suppression (74-88 Mb; Figure 4b). Fusion events might be facilitated by repeats, and in line with this we saw a distinct local increase in repeat elements near the fusion point on the great reed warbler Z chromosome (around 77.8 Mb; Figure 4b; Supplementary Figure 3).

To evaluate how the pattern of nucleotide diversity has been affected by sex-linkage, we calculated nucleotide diversity estimates based on five female and five male whole-genome resequenced great reed warblers (Supplementary Table 1b). The analysis showed a genome average nucleotide diversity of 0.0030 in both males and females (Figure 4c; Supplementary Table 12), and revealed much lower levels for both the ancestral Z and W chromosome regions (Z: 0.0007 in males and females; W: 0.0002 in females; Figure 4c; Supplementary Table 12); a pattern seen also in other bird species (reviewed in Sayres et al. 2018). Both the added-W and -Z had lower nucleotide diversity levels (W: 0.0001 in females, Z: 0.0044 in females and 0.0040 in males; Figure 4c; Supplementary Table 12) than the autosomally segregating part of chromosome 4A (0.0068 both in males and females; Figure 4c; Supplementary Table 12). The diversity of the added-Z was, however, higher than the genome average. We notice that despite its comparably recent history as sex-linked, the added-W diversity levels are similar to those of the ancestral-W.

### 2.5 Substitution rates and purifying selection among gametologs

Next, we analysed the rate of synonymous (dS) and non-synonymous substitutions (dN) of ZW gametologs to investigate whether purifying selection acts on sex-linked genes. We aligned the Z- and W-sequences of each of the 147 great reed warbler ZW gametologs (see above) together with orthologous gene copies from the zebra finch gene annotation (Z-linked genes for the ancestral region and chromosome 4A-linked genes for the added region), and calculated dS, dN and dN/dS for each pairwise comparison and gene (see Methods). Of the 41 gametologs from the ancestral sex chromosome region, 35 remained after filtering for a minimum length of 500 base pairs and a maximum dS value of 3. For the added region, 79 of the 106 gametologs remained after filtering (Supplementary Table 13).

As expected from its much older history of sex-linkage, the ancestral sex chromosome region showed higher dS and dN between the Z and W gametologs (median dS = 0.263; median dN = 0.026) compared to the added region (median dS = 0.078; median dN = 0.013) (Mann-Whitney U test; dS: *U* = 177, *p* = 1.33×10^−13^; dN: *U* = 933, *p* = 0.006; Supplementary Figure 4a,b). However, the dN/dS ratio was significantly higher for the added region (median dN/dS: 0.155; range: 0.001-0.890) than for the ancestral region (median dN/dS: 0.109; range: 0.001-0.289; Mann-Whitney U test: *U* = 1776, *p* = 0.016; Supplementary Figure 4c). This result suggests that purifying selection is generally acting on sex-linked gametologs (dN/dS < 1 for all gametologs; Supplementary Table 13), but particularly strongly so on genes being maintained both on the Z and the W chromosome over very long periods of time.

Then, we analysed substitution rates between each of the great reed warbler Z and W gametolog and the corresponding zebra finch ortholog. For gametologs on the added region, where zebra finch chromosome 4A orthologs are analysed, there was no difference between dS for W to zebra finch and dS for Z to zebra finch (Wilcoxon signed-rank test: *V =* 1699, *p* = 0.56), whereas the W gametologs showed higher dN and dN/dS to zebra finch than did Z gametologs (dN: *V* = 2582, *p* = 1.01×10^−9^; dN/dS: *V* = 2588, *p* = 5.82×10^−9^). Similarly, the genes on the ancestral sex chromosome had higher dS and dN values (Wilcoxon signed-rank test: *V =* 630, *p =* 5.82×10^−11^ in both cases) between W to zebra finch than between Z to zebra finch. Also, the dN/dS values from these comparisons differed significantly (*V* = 524, *p* = 1.11×10^−4^). These results are in line with purifying selection being less efficient on the W than the Z chromosome. Note, however, the analysis of gametologs on the ancestral region is be biased towards higher divergence values for W-linked genes as recombination suppression on the ancestral sex chromosome precedes the split between the zebra finch and great reed warbler, which makes the finch and warbler Z orthologs share more recently history.

### 2.6 Conserved and dose sensitive genes maintain W gametologs

As the W chromosome degenerates, many W gametologs are lost, and the strong signature of purifying selection on sex-linked genes (supported by the dN/dS values above) suggests that the ones being maintained on W are biased towards genes with conserved functions. To test this, we contrasted substitution rates between great reed warbler Z-linked genes and the corresponding zebra finch orthologs for (i) Z-linked genes where the W-linked gametolog has become lost, and (ii) Z-linked genes where the W-copy remains in the great reed warbler assembly. We did this for both the added and ancestral sex chromosome region, where in the latter analysis we also included 243 (non-manually curated) genes from the ancestral Z chromosome of which there was no W-copy in the gene annotation. After alignment of sequences and filtering of short (<500 bp) alignments, we analysed 273 genes from the ancestral sex chromosome (35 with and 238 without a W gene copy), and 97 genes from the added sex chromosome (79 with and 18 without a W gene copy). Z-linked genes with lost W gene copies were found distributed along the entire ancestral (Figure 5a) as well as added (Figure 5b) sex chromosome region.

**Figure 5.**
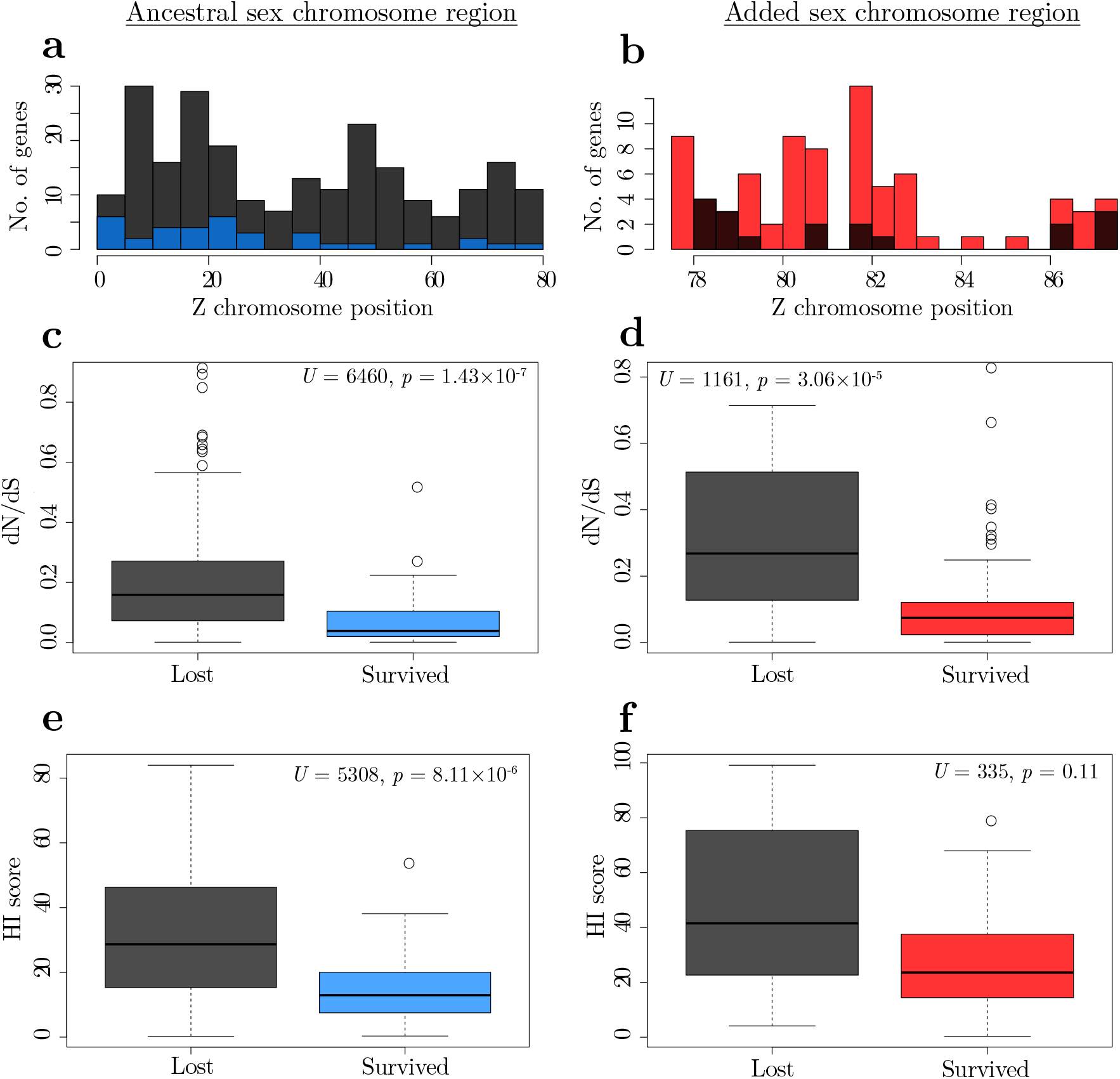
Non-random loss of W gametologs on the great reed warbler neo-sex chromosome. **(a**,**b)** Chromosome position of Z genes with lost (black) or retained W gene copy (blue/red) on (a) the ancestral and (b) added part of the sex chromosome. **(c-f)** The ratio of non-synonymous to synonymous substitution rates (dN/dS) between orthologous great reed warbler and zebra finch genes (c,d), and haploinsufficiency (HI) scores (e,f), for Z-linked genes with and without a W copy on the ancestral (c,e; blue) and the added (d,f; red) sex chromosome region. The median value is marked by the black line in each box, and the upper and lower hinges signify the first and third quartiles. The whiskers extend to no more than 1.5× of the interquartile range from each hinge.

On the ancestral part of the neo-sex chromosome, there was no difference in dS values between Z-linked genes with (median = 0.115) and without (median = 0.119) a W gene copy (Mann-Whitney U test: *U* = 4795, *p* = 0.15), while both the dN (lost W, median = 0.020; survived W, median = 0.005; Mann-Whitney U test: *U* = 6575, *p* = 3.30×10^−8^) and dN/dS (lost W, median = 0.159; survived W, median = 0.038; Mann-Whitney U test: *U* = 6460, *p* = 1.43×10^−7^) values were significantly higher for Z-linked genes without a W gene copy (Figure 5c; Supplementary Table 13). The results were similar for the added part of the sex chromosome: the dS values did not differ between Z-linked genes with (median = 0.128) or without (median = 0.130) a W gene copy (Mann-Whitney U test: *U* = 703, *p* = 0.94), while the dN values was significantly higher for Z-linked genes without a W gene copy (lost W, median = 0.031; survived W, median = 0.010; Mann-Whitney U test: *U* = 1129.5, *p* = 1.05×10^−4^), and the same was true for the dN/dS values (lost W, median = 0.268; survived W, median = 0.074; Mann-Whitney U test: *U* = 1161, *p* = 3.06×10^−5^; Figure 5d; Supplementary Table 13). The lower dN and dN/dS values for Z genes where the W-gene copy has survived supports that these sex-linked genes are under strong purifying selection for being functionally conserved.

The W chromosome is further expected to be enriched for dose sensitive genes, as haploinsufficiency (HI) will pose problems for the heterogametic sex when the W gene copy is functionally lost. We downloaded predicted HI scores based on human studies (where a lower HI value describes that a diploid gene is less able to retain its full function when a mutation disrupts one of its gene copies) from the DECIPHER database (https://decipher.sanger.ac.uk/; accessed 11 January 2019) for orthologs of our Z-linked great reed warbler genes. In line with predictions, we found for the ancestral sex chromosome region that genes with a surviving W copy had lower HI scores (*n* = 33; median HI score = 12.95) than genes that had lost their W copy (*n* = 217; median HI score = 28.66; Mann-Whitney U test: *U* = 5308, *p* = 8.11×10^−6^; Figure 5e; Supplementary Table 14). For the added sex chromosome part, a similar but non-significant pattern was found (*n* = 62 genes with a W copy: median HI score = 23.59; *n* = 8 genes without a W copy: median HI score = 41.52; Mann-Whitney U test: *U* = 335, *p* = 0.11; Figure 5f; Supplementary Table 14).

### 2.7 Evolution of sex-linked genes is highly predictable

Finally, we evaluate the consequences of sex-linkage *per se* for the evolutionary trajectory of the genes on the Sylvioidea neo-sex chromosome by making use of the rich source of available genomic data of orthologous genes in birds and other vertebrates with varying degree of sex- and autosomal linkage. From Ensembl BioMart, we downloaded dN/dS values from orthologs to sex-linked great reed warbler sex chromosome genes for one species-pair each of birds (zebra finch and chicken, *Gallus gallus*), reptiles (green anole, *Anolis carolinensis*, and bearded dragon, *Pogona vitticeps*), mammals (human, *Homo sapiens*, and house mouse, *Mus musculus*) and fish (three-spined stickleback, *Gasterosteus aculeatus*, and fugu, *Takifugu rubripes*), respectively (Supplementary Table 15). Next, we correlated these dN/dS values to each other as well as to the dN/dS values for great reed warbler gametologous gene pairs (great reed warbler Z vs. great reed warbler W) and for each great reed warbler gametolog to the zebra finch ortholog (great reed warbler Z vs. zebra finch; great reed warbler W vs. zebra finch). Interestingly, these pairwise analyses of dN/dS values showed strong positive correlations not only for comparisons within birds, but also for deeply diverged groups such as birds and fish. In fact, for comparisons involving orthologs to genes on the ancestral sex chromosome region, all correlations were positive and all except one significantly so (Spearman correlation: *p*-values < 0.05; exception: great reed warbler Z vs. great reed warbler W compared to stickleback vs. fugu; *p* = 0.051; Figure 6a), and for comparisons involving orthologs on the added region all correlations were positive and significant (Figure 6b; *p*-values < 0.05; scatter plots and *p*-values for all correlations are provided as Supplementary Figures 5 and 6).

**Figure 6.**
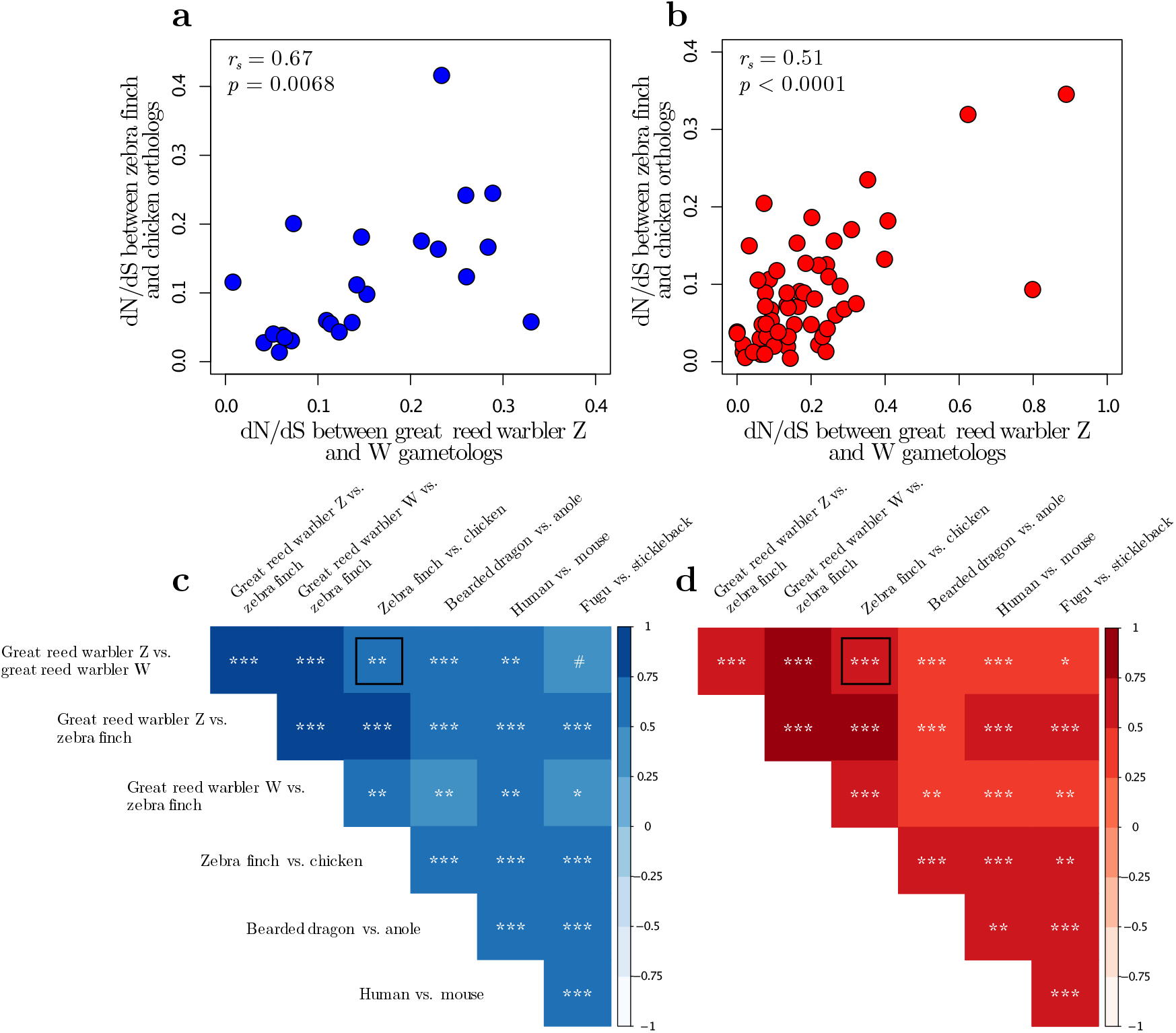
Correlated rate of evolution of orthologs in vertebrates. **(a**,**b)** Relationship between dN/dS values for great reed warbler Z and W gametologs, and dN/dS values for zebra finch and chicken Z orthologs, for genes located on (a) the ancestral and (b) the added sex chromosome region. Correlation coefficients and p-values are shown. **(c**,**d)** Correlation coefficients (heat map) and significance levels (symbols) for pairwise correlations of seven sets of dN/dS values between orthologous (or gametologous) to genes located on (c) the ancestral and (d) the added sex chromosome region. Symbols signify p-values of < 0.1 (#), < 0.05 (*), < 0.01 (**) and < 0.001 (***). The black squares mark the significance levels corresponding to the data shown in panels (a) and (b), which are highlighted as two examples of dN/dS correlations. The seven sets of dN/dS values came from comparisons of (i) great reed warbler gametologous gene pairs (Z vs. W), (ii and iii) each of the great reed warbler gametologs to zebra finch orthologs (great reed warbler Z vs. zebra finch, and great reed warbler W vs. zebra finch), and (iv-vii) orthologs for one species-pair each of birds (zebra finch and chicken), reptiles (anole and bearded dragon), mammals (human and mouse) and fish (stickleback and fugu), respectively.

## 3. DISCUSSION

Fusions between autosomes and sex chromosomes are rare in birds and have so far been reported only among Sylvioidea species (Pala et al. 2012; Sigeman et al. 2019; Dierickx et al. 2020; Leroy et al. 2019; Sigeman et al. 2020) and in the eastern yellow robin (*Eopsaltria australis*; Gan et al. 2019). Our annotated genome assembly and detailed characterisation of the Z and W chromosomes of one Sylvioidea species, the great reed warbler, provide strong evidence for translocation of a part of chromosome 4A to the ancestral sex chromosome though fusion events on both the Z and the W chromosomes. The Z fusion is covered by a single scaffold (Scaffold31) and we observe a local accumulation of repeats (LTRs) at the fusion point which may have facilitated the translocation. Synteny between the translocated, added Z chromosome region and the corresponding region on chromosome 4A in other passerines further revealed complete collinearity and maintained size (9.6 Mb), showing that no intra-chromosomal rearrangements involving this region have occurred since the fusion event. In contrast, the W chromosome, which to a large extent is covered by two sizable “superscaffolds”, is more dynamic with multiple large- and small-scaled rearrangements. Several W scaffolds contain sequences from both the ancestral and the added part, which provides strong evidence for a W fusion followed by intra-chromosomal rearrangements. A fusion event on Sylvioidea W is further supported by karyotype data in larks, family Alaudidae (Bulatova 1981; see Sigeman et al. 2019), and by sequence data in white-eyes, family Zosteropidae (Leroy et al. 2019).

The W chromosome is one of the most difficult regions in the genome to assemble due to its repetitive and haploid nature (Tomaszkiewicz et al. 2017). Karyotype data have shown that the size of the W chromosome in birds varies even over short timescales (Rutkowska et al. 2012), although much of the variation in size of W assemblies can be attributed to differences in sequencing technology, with short-read sequencing failing to scaffold repetitive regions and therefore underestimating the actual size of W (Smeds et al. 2015; Xu et al. 2019). By using long-read sequencing and optical maps we managed to assemble a total of 30.2 Mb of the W chromosome, of which 12.3 Mb could be traced back to the ancestral and 10.3 Mb to the added part. This suggests that our W assembly (considering both placed and unplaced W-linked scaffolds) approaches the size (∼21 Mb) of the latest version of the zebra finch W chromosome, which does not include the added-W (NCBI Annotation Release 105: bTaeGut2.pat.W.v2; Rhie et al. 2020). Despite these difficulties associated with assembling W chromosomes, it seems clear that the ancestral-W is much smaller than the ancestral-Z chromosome (∼80 Mb) in most birds, including the great reed warbler. In contrast, and despite a few Mb-large deletions of gene poor regions, our assembly of the added-W region (10.3 Mb) is longer than the added-Z region (9.6 Mb), which is likely the result of repeat accumulation. A relative increase in size of the sex-limited chromosome (W/Y) compared to its chromosome copy (X/Z) has been observed in other lineages as well (e.g. *Drosophila miranda*; Bachtrog et al. 2019).

In present day great reed warblers, recombination is suppressed across the entire added sex chromosome region, meaning that recombination stopped between the fusion event and today. We show that the added region continued to recombine for several Myr after the fusion event and thus acted as a second PAR in the ancestors of the great reed warbler. The relative importance of large-scale rearrangement events (e.g. large inversions) compared to more gradual and smaller mutations for the evolution of recombination suppression has been debated (Ponnikas et al. 2018). Up until recently, most evidence of large inversions for causing recombination cessation had come from old and heavily differentiated sex chromosome systems such as from birds and mammals. Empirical data from more recently formed sex chromosome systems has however brought the insight that both inversions and gradual events can lead to recombination suppression (reviewed in Wright et al. 2016). The detected rearrangements in the added W region in our data (there were no signs of rearrangements on the added Z chromosome) may have contributed mechanistically to recombination suppression. However, rearrangements are also more likely to become fixed in regions of already low recombination, making the distinction between cause and consequence of suppression extremely difficult. Our data show that recombination suppression in great reed warblers seems to have progressed in a non-linear, mosaic, and small-scaled manner along the added part of the neo-sex chromosome. This does not follow an expected pattern of large-scale rearrangements starting from the fusion point. Instead, it is in line with a local recombination suppression process, possibly mechanistically driven by a combination of small inversions, repeat accumulation and heterochromatinization (cf. Ponnikas et al. 2018). This mosaic pattern makes it difficult to define evolutionary strata on the added region of the neo-sex chromosome, which contrasts the situation on the ancestral avian sex chromosomes (Zhou et al. 2014; Xu et al. 2019).

As expected, we observe clear consequences of sex-linkage and recombination suppression on the Sylvioidea neo-sex chromosomes, and in particular so for the sex-limited copy (W) which does not recombine in either sex. Both the ancestral and added W chromosome regions showed pronounced accumulation of repeat elements (64% and 35%, respectively; mainly LTRs) and low nucleotide diversity (0.00024 and 0.00014, respectively). The repeat content of the great reed warbler ancestral-W scaffolds (64%) was higher than in the short-read sequenced collared flycatcher W chromosome (FicAlb1.5: 49%; Smeds et al. 2015) and slightly lower than the W chromosome in the long-read sequenced chicken genome (galGal5: ∼68%; Kapusta & Suh 2017). The Z chromosome had similar repeat levels as autosomes, but we found support for reduced nucleotide diversity on the ancestral-Z (0.00074) compared to the genome-wide average (0.00301), and on the added-Z (0.00442) compared to the autosomal part of chromosome 4A (0.00678). In addition of supporting consequences of sex-linkage also on avian Z chromosomes, this result highlights the importance of comparing chromosomes with similar properties (e.g. size and gene density) in intra-specific analyses, and ideally homologous chromosomes in inter-specific analyses, when evaluating the genomic consequences of sex-linkage (Julien et al. 2012). Lower nucleotide diversity on sex chromosomes have been observed in many lineages (Sayres 2018), but the relative importance of effective population size effects and selection is poorly understood. In addition to this, the female-specific inheritance of the W chromosome is also expected to contribute to lower nucleotide diversity than on the Z chromosome and autosomes, as birds like e.g. mammals have male-biased mutation rates (Ellegren & Fridolfsson 1997). Regardless of the cause, the extremely low nucleotide diversity on the ancestral as well as added W chromosome regions likely diminishes the evolutionary potential of the neo-sex chromosome W copy.

Instead, W-linked genes are likely to either become lost by degeneration and drift, or be preserved through purifying selection. Our analyses of synonymous and non-synonymous substitutions, and haploinsufficiency, strongly suggest that the W chromosome is enriched for dose sensitive and conserved genes that are being maintained by purifying selection. The dN/dS ratio between gametologs was low for the added region and even more so for the ancestral region, which can be explained by particularly strong purifying selection on the few surviving genes on the ancestral W chromosome region. Together these results suggest that W gametologs with conserved functions are being maintained functionally by purifying selection over long evolutionary timescales, and that the new set of sex-linked genes on the added part of the Sylvioidea neo-sex chromosomes mimics the ancestral avian sex chromosome at an earlier stage of its evolution. Our set of 41 gametologous gene pairs on the ancestral W region was highly overlapping with the gene set found in the flycatcher W chromosome (that is entirely ancestral; Smeds et al. 2015), which shows that the ancestral W chromosome in the great reed warbler has not undergone more pronounced degeneration compared to other songbirds. We also observe that Z-linked genes with a remaining copy on the W chromosome are more conserved and dose sensitive than genes whose copy has been lost from the W chromosome, a pattern that is concordant with studies of ZW-gene pairs in chicken (Bellott et al. 2017). In general, although degeneration seems a likely long-term evolutionary trajectory for a majority of W-linked genes, apparently some W genes are maintained for long periods of time by purifying selection. We did not find any genes on the added-W without gametologs on the added-Z. This is in contrast to, e.g., the situation in some mammals, where a few genes have been added to the Y chromosome (Cortez et al. 2014) and in *Drosophila miranda*, where intense gene translocation to the Y chromosome has occurred (Bachtrog et al. 2019).

Strong purifying selection on sex-linked genes are thought to drive patterns of convergent sex chromosome evolution across different taxonomic groups with independently evolved sex chromosome systems, such as birds (ZW) and mammals (XY) (Bellott et al. 2017). Our results extend these conclusions by showing that gene rate evolution (dN/dS) is strongly positively correlated between widely diverged taxonomic groups, regardless of whether the genes are autosomal or sex-linked, Z- or W-linked, or located on newer or older parts of the sex chromosome. For example, we find highly significant correlation between dN/dS-values for a set of great reed warbler Z and W gametologs, and dN/dS for orthologs to these genes in two lizard species. The majority (99.8%) of dN/dS-values are < 1, and all pairwise correlations are positive and significant (except for one with p = 0.051). This strongly suggests that these broad taxonomic trends are driven by different levels of purifying selection acting on genes with more or less conserved essential functions on a deep phylogenetic level. Still, we cannot exclude the action of correlated positive selection although we believe it has a minor influence, especially for W-linked genes. We conclude that the highly predictable evolutionary trajectory of sex-linked genes in both birds and mammals (cf. Bellott et al. 2017) is driven partly by sex-linkage *per se* (e.g. due to small effective population size and inefficient selection), partly by different degrees of functional conservation of specific genes.

## 4. MATERIALS AND METHODS

### 1.1 DNA extraction

High molecular weight DNA was extracted from the blood of a juvenile female great reed warbler (used to produce the genome assembly; Supplementary Table 1a) and kept in -80°C in SET buffer (0.15 M NaCl, 0.05 M Tris, 0.001 M EDTA, pH 8.0) using standard phenol-chloroform extraction (Sambrook et al. 1989) with initial RNase treatment and Proteinase K digestion and final collection of purified DNA on a glass rod.

### 1.2 RNA extraction

RNA from the genome individual was extracted from snap frozen liver, heart and muscle tissue that was kept in -80°C. The extraction was carried out using RNeasy mini kit (Qiagen, cat no. 74104), with 15 min on-column DNase treatment, according to the manufacturer’s instructions. The RNA was used for generating short-read Illumina RNA-seq (Supplementary Table 1a).

### 1.3 Iso-Seq libraries

To construct cDNA libraries for PacBio sequencing (Iso-seq), mRNA was first purified from total RNA from the genome individual by two rounds of polyA-selection using Poly(A)Purist MAG kit (Ambion, cat nr AM1922) according to the manufacturer’s instructions. The mRNA was used for cDNA synthesis according to “Procedure and Checklist – Isoform Sequencing (Iso-Seq) Using the Clontech SMARTER PCR cDNA Synthesis kit and BluePippin Size-selection System” (Pacific Biosciences) and was prepared to an Iso-Seq library according to “Guidelines for Preparing cDNA Libraries for Isoform Sequencing (Iso-Seq) User Bulletin” (Pacific Biosciences).

### 2.1 Assembly strategy

#### 2.2.1 Long-read *de novo* assembly

From 108 PacBio SMRT cells (RSII chemistry P6-C4), we sequenced 13,173,355 subreads (117405507246 bases in total) with a mean length of 8,912 bp and a N50 read length of 11,757 bp (Supplementary Table 1a). The genome coverage of the error corrected reads, considering only reads >10 kb, was 24.9×. FALCON (Chin et al. 2016) assembler was used to make a draft assembly (consisting of primary and associated contigs representing structural variants) based on the all subreads longer than 8 kb (configuration file provided in Supplementary Methods). The draft assembly was error corrected twice using Quiver (Chin et al. 2013).

#### 2.2.2 Error correction using Illumina data

The draft assembly was error corrected again using Illumina paired-end reads (2×150 bp) from the same individual that was used for PacBio sequencing (Supplementary Table 1a). The paired-end reads were first trimmed using Trimmomatic v.0.36 (Bolger et al. 2014) using the following settings: TruSeq3-PE.fa:2:30:10 LEADING:15 TRAILING:30 SLIDINGWINDOW:4:20 MINLEN:90. Of the original 479,067,329 read pairs, 356,222,474 (74.36 %) survived trimming of both reads. The surviving read pairs were aligned to the PacBio draft assembly using bwa mem v.0.7.15 (Li & Durbin 2009), transformed to bam format and sorted with samtools v.0.1.19 (Li et al. 2009). Duplicate reads were then removed using picard MarkDuplicates v.2.0.1 (http://broadinstitute.github.io/picard). The number of aligned and deduplicated reads were 632,245,634. The aligned reads were then used to polish the genome assembly with Pilon version 1.17 (Walker et al. 2014) using options --genome --frags -diploid --fixbases. The error corrected draft assembly consisted of 8,274 contigs in total, with a total genome length of 1.35 Gb. From this draft assembly, we extracted only the primary contigs (5,419 contigs with a total genome size of 1.22 Gb and N50 of 3.7 Mp).

#### 2.2.3 Misassembly detection and scaffolding with linked-read data

We used Chromium linked-read data (10x Genomics) to identify and break apart scaffolds at suspected misassembled sites, and then to increase contiguity through scaffolding (Supplementary Table 1a). The linked-read data was demultiplexed and transformed to fastq format using the supernova mkfastq program from Supernova v.1.1.5 (Weisenfeld et al. 2017). The barcodes from the fastq data were processed and error corrected using the longranger basic (v.2.1.6) tool from 10x Genomics. After processing, 408 million read pairs remained with 95.6% whitelisted barcodes and a barcode diversity of 752,722. We used tigmint (Jackman et al. 2018) with settings as=100, depth_treshold=65, minsize=2000, number of mismatches=5 to break contigs at suspected misassemblies and low-quality regions. Next, arcs (Yeo et al. 2017) was used to scaffold the contigs using settings c = 5, e=30000 and r=0.05, followed by links v1.8.5 (also from the arcs pipeline) using settings -a 0.9 and -l 5. The resulting draft assembly consisted of 9,823 scaffolds (of which 6,727 are longer than 1,000 bp). The total length was still 1.2 Gb, scaffold N50 was 19Mb and GC 43 %.

In order to detect additional misassemblies in the polished and scaffolded draft assembly, we aligned it to the genome assembly of the zebra finch (taeGut3.2.4; Warren et al. 2010, downloaded from Ensembl; Yates et al. 2020) using SatsumaSynteny v2.0 (Grabherr et al. 2010). For each chromosome in the zebra finch assembly, we extracted genomic coordinates from great reed warbler draft assembly scaffolds that aligned to this chromosome using BEDTools merge (v2.27.1; Quinlan and Hall 2010) with option -d 100000. Only alignments to the zebra finch genome that were larger than 100 kb were considered. Then, we used BEDTools complement to create genomic ranges also for the genomic regions that were not part of the zebra finch alignment dataset, if these ranges were longer than 100 bp. We then used BEDTools getfasta to make a new fasta file where scaffolds were cut according to these genomic ranges. This means that scaffolds that align to two separate zebra finch chromosomes (or only partly to a zebra finch chromosome) will be split into two in the breakpoint region. This new fasta file was processed with arcs and links for a second round of scaffolding using the same settings as above. The new draft assembly consists of 7,985 scaffolds (of which 6,531 are longer than 1,000 bp). While the length of the assembly remained almost unchanged, this scaffolding step increased the scaffold N50 to 21 Mb.

#### 2.2.4 Scaffolding with optical mapping data

We then used Bionano optical mapping data to further increase the contiguity of our data (Supplementary Table 1a). DNA from the same individual that was used for the reference genome was extracted using the agarose plug method from blood (in SET buffer). Two enzymes were used; BSPQI and BSSSI. The data from each enzyme was first assembled into separate *de novo* assemblies (using the script pipelineCL.py from Bionano Solve with settings -U -d -T 228 -j 228 -N 4 -i 5). The script runTGH.R from Bionano Solve (version Solve3.1_08232017) was used to anchor the scaffolds from the draft assembly to the optical mapping assemblies using standard settings, options: -e1 BSPQI -e2 BSSSI and using the provided configuration file ‘hybridScaffold_two_enzymes.xml’. Scaffolds with a combined length of 1.1 Gb (N50: 19 Mb) were anchored in the new hybrid assembly, which had an N50 value of 20.5 Mb.

#### 2.2.5 Gapfilling and additional error correction

To fill in gaps in the draft assembly, we used gapfiller (Nadalin et al. 2012) and aligned PacBio reads. After gapfilling, the draft assembly was error corrected once using PBjelly (English et al. 2012) and twice using quiver. All scaffolds shorter than 1,000 bp were removed from the assembly.

#### 2.2.6 Splitting up of chimeric scaffolds

Seven scaffolds were manually broken apart at misassembled sites, identified through (i) the linkage map data (Ponnikas et al. in prep.), (ii) aligned genomic Illumina reads from the same individual that was used to create the reference genome, and (iii) synteny information from the zebra finch genome. The scaffolds were split using the script https://github.com/NBISweden/NBIS-UtilityCode/SplitFastaAndGFF.cc.

#### 2.2.7 Removal of redundant scaffolds

We removed redundant scaffolds that represent haplotypes of another scaffold (“haplotigs”) by using the purge haplotigs pipeline (Roach et al. 2018). For the pipeline we first estimated coverage for each scaffold by mapping PacBio subreads to the assembly using minimap2 v.2.13 (Li 2018). Based on the coverage distribution in the genome we set 60x as the threshold between haploid and diploid coverage. Any scaffold with a diploid coverage less than 80% was considered as a suspect haplotig and was searched against other scaffolds within the software. We removed scaffolds that had a best match coverage of at least 95% to its best hit. This resulted in the removal of 3,468 scaffolds with a mean length of 14,543 bp (range: 1,001 bp – 920,475 bp).

#### 2.2.7 Repeat masking

We *de novo* predicted repeats in the assembly using RepeatModeler v.1.0.8 (Smit & Hubley 2008-2015) with the option -engine ncbi. We then ran RepeatMasker v4.0.7 (Smit et al. 2013-2015) with the output from RepeatModeler and a custom library with manually curated repeat elements from different bird genomes provided by Alexander Suh (Uppsala University; fAlb15_rm3.0_aves_hc.lib) with options -a -xsmall -gccalc.

#### 2.2.8 Cross-species synteny visualization

To produce the circos plot (Figure 1), we ran SatsumaSynteny v2.0 (Grabherr et al. 2010) between the great reed warbler genome and the great tit genome (Parus_major1.1; GCA_001522545.2; Laine et al. 2016, downloaded from Ensembl). We then calculated the length (bp) of all matches between great reed warbler scaffolds and great tit chromosomes. Scaffolds that matched with more than 100 kb and more than 1% of its length to any great tit chromosome were kept for visualization.

This left 123 scaffolds with a total length of 1.10 Gb. We then used the tool bundlelinks (from circos-tools-0.23; Krzywinski et al. 2009) and settings -max_gap 100000 -strict -links to produce larger ranges of matches between the species. Lastly, we removed scaffolds that had been identified as W-linked (see above), leaving 100 scaffolds with a total length of 1.07 Gb. Alignments between great reed warbler scaffolds and great tit chromosomes were plotted in circos v.0.69-8 (Krzywinski et al. 2009).

### 2.2 Annotation

#### 2.2.1 RNA sequencing data generation and processing

We trimmed RNA-seq Illumina reads (Supplementary Table 1a) with trimmomatic v 0.36 (Bolger et al. 2014) using the parameters TruSeq3-PE-2.fa:2:30:10 SLIDINGWINDOW:4:5 LEADING:5 TRAILING:5 MINLEN:25, as suggested in the Trinity (Grabherr et al. 2011) documentation. Of 512,409,808 raw Illumina reads, 507,717,996 reads remained after trimming. For a reference-guided transcriptome assembly, the draft genome assembly was indexed using bowtie2 v.2.3.2 (Langmead & Salzberg 2012). We then aligned the trimmed RNA-seq reads separately for each tissue to the assembly using tophat v.2.1.1 (Kim et al. 2013) with the option --library-type=fr-firststrand. Accepted hits from the three tissues were merged into one file using samtools (v.1.3; Li et al. 2009) merge. The reads were then assembled using StringTie v.1.3.3 (Pertea et al. 2015) and cufflinks v.2.2.1 (Trapnell et al. 2010). The output was transformed from GTF to GFF format using gffread (Pertea & Pertea 2020) with option -E.

We performed a *de novo* assembly using the trimmed RNA-seq Illumina reads combined for all three tissue samples in Trinity v.2.3.2 (Grabherr et al. 2011) with the options “--seqType fq -- SS_lib_type RF”.

#### 2.2.2 Gene builds

We then predicted gene models using MAKER v.3.00.0 (Holt et al. 2011; Campbell et al. 2014). The reference-guided and *de novo* RNA-seq assemblies from Illumina short read data (Supplementary Table 1a) were provided to MAKER along with an assembly of the Iso-seq data as species-specific evidence. As additional evidence manually reviewed protein sequences (556,825 proteins) from the Swissprot section of the Uniprot database were downloaded (2018-03) (Magrane & Consortium 2011), along with protein files from chicken (Gallus_gallus.Gallus_gallus-5.0.pep.all.fa; containing 30,252 proteins) and zebra finch (Taeniopygia_guttata.taeGut3.2.4.pep.all.fa; containing 18,204 proteins). We provided the repeat library fAlb15_rm3.0_aves_hc.lib and the output from RepeatMasker (see above) as input for repeat masking with RepeatMasker and RepeatRunner (Yandell 2006) which are run internally by MAKER. We used the *ab-initio* gene finder Augustus v.3.2.3 (Stanke et al. 2006) with the pre-trained profile of chicken. Gene builds were constructed in MAKER, using 1) only the extrinsic evidence (proteins and transcripts), and 2) combining the gene builds from extrinsic evidence sequences with *ab-initio* predictions in Augustus. As the evidence run performed better than the *ab-initio* run (evaluated using BUSCO: 82.5% complete genes compared to 66.6%; and through visual inspection), we used the evidence run as a base and complemented it with the Augustus annotation track created during the *ab-initio* run using an in-house perl script (https://github.com/NBISweden/AGAT/bin/agat_sp_complement_annotations.pl). Another in-house perl script (https://github.com/NBISweden/AGAT/bin/agat_sp_fix_longest_ORF.pl) was used to improve the ORF start and end positions, to improve fragmented and missing genes.

#### 2.2.3 Functional annotation

We inferred the function of genes and transcripts using the translated CDS features of each coding transcript. To retrieve a gene name and function of gene, we (i) blasted the predicted protein sequence of each transcript against the Uniprot/Swissprot reference dataset and (ii) ran the same sequences in InterProscan v-5.7-48 (Jones et al. 2014). Then, the Annie annotation tool (Tate et al. 2014) was used to extract relevant metadata into predictions for canonical protein names and functional predictions. This resulted in 20,807 gene models with 1,717 gene models without functional annotations. Gene names were inferred with a best blast hit approach using the Uniprot/Swissprot reference dataset. In total, 18,559 genes were named, of which 2,312 had duplicate gene names.

We predicted tRNA using tRNAscan v.1.3.1 (Lowe et al. 1997) (450 tRNAs) and other ncRNAs using the RNA family database Rfam v.11 (Nawrocki et al. 2014).

A lift-over annotation to the great reed warbler genome was done using the (i) zebra finch (Taeniopygia_guttata.taeGut3.2.4.94; 17,487 genes) and (ii) chicken (Gallus_gallus.Gallus_gallus-5.0.94; 18,345 genes) ensemble gene annotations. To generate a lift-over, we first did pairwise alignments between the great reed warbler genome and the other genomes using SatsumaSynteny v.3.0. Then, Kraken (Zamani et al. 2014) was used to project the annotations from one genome to another using the pairwise alignments. Finally, an in-house script (https://github.com/NBISweden/AGATagat_sp_kraken_assess_liftover.pl) was used to handle the gene lift-overs. From the zebra finch, 14,686 genes were successfully lifted over (642 genes mapping to several locations), and 14,466 from chicken (571 genes to several locations).

OrthoMCL v2.0.9 (Li et al. 2003) was used to group orthologous protein sequences. We used proteins downloaded from Ensembl of mouse (*M. musculus*; Mus_musculus.GRCm38), anole lizard (*A. carolinensis*; Anolis_carolinensis.AnoCar2.0), human (*H. sapiens*; Homo_sapiens.GRCh38), chicken (*G. gallus*; Gallus_gallus.Gallus_gallus-5.0), tammar wallaby (*N. eugenii*; Notamacropus_eugenii.Meug_1.0), gray short-tailed opossum (*M. domestica*; Monodelphis_domestica.monDom5) and a set of protein of zebra finch (*T. guttata*; Taeniopygia_guttata.taeGut3.2.4). In total, 7,623 OG were common to all species, and 374 OG are specific to *A. arundinaceus*.

#### 2.2.4 Genome assembly of second great reed warbler female

To verify the correctness of some W-linked scaffolds, a genome assembly from linked reads sequenced from the blood of another great reed warbler female (Supplementary Table 1b) was assembled using supernova run 2.1.0 (Weisenfeld et al. 2017). Genome assembly statistics are provided in Supplementary Table 2 and 3.

### 2.3 Sex chromosome analyses

#### 2.3.1 Finding sex-linked scaffolds

We aligned paired-end sequence data (Illumina HiseqX 150 PE) from five female and five male great reed warbler individuals to the reference genome (sample information in Supplementary Table 1b). The reads were trimmed using Trimmomatic v.0.36 (Bolger et al. 2014) prior to alignment using settings TruSeq3-PE.fa:2:30:10 LEADING:15 TRAILING:30 SLIDINGWINDOW:4:20 MINLEN:90. Reads were aligned using bwa mem v.0.7.15 (Li & Durbin 2009) (option -M), alignments were sorted and converted to the bam format using samtools v.1.7 (Li et al. 2009) and reads were deduplicated with picard MarkDuplicates v.2.0.1 (http://broadinstitute.github.io/picard). We followed the general method from Smeds et al. (2015) for identifying W-linked scaffolds by first parsing the alignment files for reads with any mismatching base pairs (bam file tag NM:i:0). Then, per site genome coverage was calculated using samtools depth for reads with a minimum mapping quality of 20 and a maximum read depth of 80 (in order to avoid genome coverage values from repetitive regions). All genome coverage values were normalized between samples based on the total number of reads in the trimmed fastq files. The normalized coverage values were summed for each sex (5 females and 5 males), and the per-sex median coverage for each scaffold was calculated. We considered scaffolds where the male coverage was zero while the female coverage was > 25X to be W-linked. This cut-off resulted in 50 W-linked scaffolds with a median female coverage of 68.03x (mean coverage 61.83), and a mean length of 605 kb (median 58 kb). Of these scaffolds, 15 were represented in the gene annotation. The gene annotations from these 15 scaffolds were manually curated and grouped as “W-scaffolds”. The 35 scaffolds not present in the annotation file were grouped as “random W-scaffolds” (Supplementary Table 6,7).

To identify Z-linked scaffolds, we utilized the difference between the median coverage values for males and females (following the same method as above) but also the difference in heterozygosity. As females are haploid for Z-linked scaffolds while males are diploid, we expect them to differ in this measurement. We calculated inbreeding coefficients (F) for each scaffold using vcftools v0.1.15 (Danecek et al. 2011) with option --het and calculated the median for each sex. Next, we filtered the scaffolds for Z-linkage based on two criteria; a) either the median coverage in females were less than 55% of the male coverage, or b) the median female coverage was less than 65% and the heterozygosity value for males and females had an absolute difference of more than 0.1. Using this method, 22 scaffolds were considered to be Z-linked. The mean length of these were 4.03 Mb (median 31 kb). Of these 22 scaffolds, 8 were represented in the gene annotation file. Same as with the W-linked scaffolds, we considered these 8 to be the “Z-linked” scaffolds and the other ones to be the “random Z-linked scaffolds” (Supplementary Table 6,7). A linkage map analysis (Ponnikas et al. 2020) using a pedigree of 511 great reed warblers assigned seven of these eight Z-linked scaffolds to the same linkage group. The one scaffold that was not assigned (Scaffold492) was relatively short (0.6 Mb) and had no RADseq SNPs used in the linkage mapping. An additional scaffold was identified through the linkage map analysis as belonging to the Z chromosome: Scaffold92. Six of these sex-linked scaffolds (Supplementary Table 7) could be anchored (i.e., ordered and oriented) successfully in the Z linkage group (Ponnikas et al. 2020). And lastly, Scaffold217 was identified as the pseudoautosomal region (PAR) according to the linkage map. This scaffold is 0.9 Mb in length, contains the PAR genes identified in other songbird species and had equal coverage values between the female and male great reed warblers (Ponnikas et al. 2020).

#### 2.3.2 Finding chromosome matches of sex-linked scaffolds

For comparisons between the ancestral to the added sex chromosome region on a DNA level, we classified those sex-linked scaffolds that were represented in the gene annotation as either ancestral or added using whole-genome alignments to the zebra finch and the collared flycatcher. In these species, the added sex-linked region would correspond to chromosome 4A and the ancestral sex chromosome to chromosome Z. To get reliable genomic positions for the great reed warbler sex-linked scaffolds we used the following method: We extracted all genomic ranges where a sex-linked scaffold (containing at least one gene in the annotation) had aligned to both the zebra finch and flycatcher genome. Among these ranges, we considered a genomic region as belonging to the ancestral sex chromosome (Z) if the same range aligned to chromosome Z (or Z_random) in both the zebra finch and flycatcher. In addition, genomic ranges that aligned in one species to chromosome Z (or Z_random) and an unplaced scaffold in the other species was accepted. The same was done for the added sex chromosome region (i.e. alignments to chromosome 4A or 4A_random). Genomic ranges shorter than 10 kb were ignored. This resulted in genomic ranges across 8 Z-linked scaffolds with a combined length of 87.2 Mb, and 11 W-linked scaffolds with a combined length of 22.4 Mb (Supplementary Table 6a,b).

### 2.3.3 Finding orthologs and manual curations of sex-linked genes

The different gene builds generated in MAKER were imported into WebApollo (Lee et al. 2013) where we manually curated 172 gametologous gene pairs. The identification of gametologs was done in the following way: We went through all W-linked scaffolds with gene annotations (i.e. the 15 scaffolds mentioned above). Each gene was blasted (blastx) against the non-redundant protein sequence database on NCBI and manually curated and the best supported isoform for each gene was selected. The same was done for the scaffolds marked as “Z-linked” and aligning either fully or partly to chromosome 4A in the zebra finch genome (i.e. two scaffolds). To identify gametologous gene pairs on the added sex chromosome region (chromosome 4A) the W-linked and Z-linked genes needed to fulfil two of the following three criteria; i) the gene should be flanked by the same genes as in the zebra finch (or the chicken if the gene was not placed within the zebra finch genome (i.e on a random chromosome or on Un), ii) there should be lift-over evidence to the same transcripts in either zebra finch or chicken and iii) the genes should belong to the same orthology group based on an orthology analysis done using orthoMCL.

For the ancestral sex chromosome, the gene order for the W-linked genes is expected to be heavily scrambled compared to the ancestral gene order. Therefore, we only accepted W-linked genes that had orthology evidence from the OrthoMCL analysis as well as from the gene annotation lift-over analyses to a certain gene. For all of those gene transcripts, we searched for transcripts present in the orthology analysis that were located on a Z-linked scaffold. These Z-linked scaffolds were accepted based on the same criteria as the 4A-linked genes (i.e. also having either gene order or liftover support). Z-linked copies of four accepted W-linked genes were missing from the orthology analysis, but were found in the gene annotation as they were located between the expected genes (i.e. conserved gene order) in the zebra finch or chicken, and had lift-over evidence matching the same gene as the W-linked gene copy. We identified 41 gametologous gene pairs from the ancestral sex chromosome with these criteria.

In total, we found 131 genes belonging to the added sex chromosome region (homologous to chromosome 4A or 4A_random in zebra finch). Two genes were placed on zebra finch chromosome 4A_random, but in the correct place according to synteny in chicken. Of the 131 genes on the added sex chromosome region, we found 106 gametologous gene pairs, i.e. where both the Z-transcript and W-transcript were identified according to the criteria described above. From the remaining 25 genes, three genes had a W copy but insufficient ortholog evidence and for 22 genes we found only Z-linked transcripts.

We also identified 277 Z-linked genes without a W-copy as follows: First, we downloaded information on orthologs from the following species: anole lizard (*Anolis carolinensis*; AnoCar2.0; GCA_000090745.1), emu (*Dromaius novaehollandiae*; droNov1; GCA_003342905.1), great spotted kiwi (*Apteryx haastii*; aptHaa1; GCA_003342985.1), chicken (*Gallus gallus*; GRCg6a; GCA_000002315.5), collared flycatcher (*Ficedula albicollis*; FicAlb_1.4; GCA_000247815.1), budgerigar (*Melopsittacus undulatus*; Melopsittacus_undulatus_6.3; GCA_000238935.1), great tit (*Parus major*; Parus_major1.1; GCA_001522545.2), blue-crowned manakin (*Lepidothrix coronata*; Lepidothrix_coronata-1.0; GCA_001604755.1) and duck (*Anas platyrhynchos platyrhynchos*; CAU_duck1.0; GCA_002743455.1). From these, we extracted only the genes that were present in all species and that were one-to-one orthologs in all species. Then, we selected only the genes that grouped with a single great reed warbler transcript in the ortholog analysis, and lastly, we selected only those transcripts corresponding to zebra finch transcripts located on either the Z chromosome or Z_random. Two of these transcripts were also present in our gametolog analysis, meaning that they have a W copy (corresponding to zebra finch Ensembl gene IDs ENSTGUT00000000103 and ENSTGUT00000001787) so these were removed. We extracted these great reed warbler transcripts (172 manually curated ZW gene pairs and 277 uncurated Z sequences) from the reference genome and added the zebra finch transcript for each gene. The sequences were aligned using the codon-aware aligner prank v.170427 (Löytynoja 2014) and removed gaps using gblocks v.0.91b (Castresana 2000). After filtering for a minimum length of 500 base pairs and dS lower than 3, 79 added sex chromosome gene pairs remained, and 18 Z-linked genes from the added region where the W-linked gene copy had disappeared. On the ancestral sex chromosome, 35 gene pairs remained after filtering. Of the uncurated ancestral Z-linked genes without a W copy, 238 remained after filtering. We calculated pairwise substitution rates between the three sequences (great reed warbler Z and W, and zebra finch) per gene using codeml from the PAML package v4.9 (Yang et al. 2007).

#### 2.3.3.4 Gametolog extraction

For all gametologous gene pairs (*n* = 172), we *in silico* extracted Z-sequences from males of five Sylvioidea species (clamorous reed warbler, *Acrocephalus stentoreus*; marsh warbler, *A. palustris;* western olivaceous warbler, *Iduna opaca*; Savi’s warbler, *Locustella luscinioides* and bearded reedling, *Panurus biarmicus*; Supplementary Table 1b) based on aligned reads to a version of the great reed warbler reference genome which had been parsed for all W-linked scaffolds (*n* = 50). The reason behind this approach was to avoid unequal mapping success to the Z-versus W-scaffolds due to recombination suppression being younger than the speciation dates. The sequences were obtained by extracting the major allele from either one or two males of the same species (depending on the available data). We also added one-to-one orthologs to these genes from the following seven species as outgroups: two non-Sylvioidea oscine passerines (great tit, *Parus major*, and zebra finch, *Taeniopygia guttata*), one suboscine passerine (blue-crowned manakin, *Lepidothrix coronata*), budgerigar (*Melopsittacus undulatus*), one Galloanserae (*Gallus gallus*), one Palaeognathae (emu, *Dromaius novaehollandiae*) and green anole (*Anolis carolinensis*). The ortholog information was downloaded from BioMart along with the CDS sequences from these genes.

#### 2.3.3.5 Finding orthologs of autosomal genes

To get a set of autosomal genes, we downloaded one-to-one orthologs from the same set of outgroup species (*n* = 7; see above) which were not on chromosome 4A or Z in the zebra finch genome (i.e. on chromosomes 1, 1A, 1B, 2-15, or 17-28) based on gene information on BioMart (accessed on 30 May 2019). Using the zebra finch transcript ID, we then searched for these genes in the orthoMCl group.txt file and selected those genes where there was only a single great reed warbler transcript in the ortholog file. Using this method, which resulted in 3,570 genes, we wanted to ensure that we were using only single copy orthologs. We then selected 100 random genes (using the bash command shuf -n 100) and extracted autosomal sequences from the different Sylvioidea species using the positions of these transcripts in the gene annotation.

#### 2.3.3.6 Recombination suppression analyses

To evaluate the timing of recombination suppression for each great reed warbler gametolog, we searched for the most likely phylogenetic position of the great reed warbler W gametolog on a fixed and dated phylogeny of the six Sylvioidea species and seven outgroup species (consisting of either Z-linked or autosomal sequences; see above). The phylogenetic position of each W gametolog was estimated by applying the evolutionary placement algorithm (EPA), so that the branching position of the W on the fixed phylogeny would represent the time since divergence to its Z gametolog, and therefore the timing of recombination suppression between Z and W. The topology of this phylogeny was compiled from Jarvis et al. (2014) and Oliveros et al. (2019). This fixed topology was then dated with MCMCTree in PAML v.4.8a (Yang et al. 2007) using alignments of 69 concatenated autosomal sequences from all 13 species (see above) and the following calibration times (also from Jarvis et al. 2014 and Oliveros et al. 2019): divergence of lizard-bird (255.9 - 299.8 Myr), Palaeognathae-Neognathae (66 - 99.6 Myr), Psittaciformes-Passeriformes (51.81 - 66.5 Myr) and Suboscines-Oscines (27.25 - 56 Myr; Supplementary Figure 2).

Sequence alignments for the each gametolog (including the Z and W sequence from the great reed warbler, in addition to sequences from five other Sylvioidea species and seven non-Sylvioidea species; see above) were constructed with PRANK v.170427 (Löytynoja 2014). After trimming by Gblocks v.0.91b (Castresana 2000) and filtering for short alignments (> 500 bp), 24 genes from the ancestral sex chromosome and 49 genes from the added sex chromosome region remained (note, that these 49 genes are in fact autosomal in all non-Sylvioidea species). We then separated the W sequence from each alignment. We estimated the evolution model parameters for each gene alignment on the fixed topology with the parallel version of RAxML v.8.2.12 (raxmlHPC; Stamatakis et al. 2014) using the following options: -m GTRGAMMAX -f e. Next, we estimated the likelihood weight ratio (LWR) for each possible phylogenetic position of the great reed warbler W gametolog on the fixed topology with EPA-ng (Barbera et al. 2019), using the model parameters file from the RAxML (for --model), the gene alignment file (for --ref-msa), the fixed topology file (for --tree), the number of tested branches in the fixed topology file (for --filter-max) and the great reed warbler W sequence (for --query), together with the following options: --no-heur --raxml-blo --filter-min-lwr 0. For the genes on the ancestral sex chromosome we tested all phylogenetic positions, whereas for the genes on the added part we tested only positions on the gene tree comprising all six Sylvioidea species and an outgroup great tit. We considered EPA placements on all internal branches and the edge of great reed warbler (LWR-values for adjacent edges were added to the LWR of the closest internal branch or the great reed warbler edge).

### 2.3.4 Population genomics

We used freebayes version 1.1.0 (Garrison & Marth 2012) to call genotypes in the whole-genome resequenced five males and five females great reed warblers mentioned above (Supplementary Table 1b). For the females, we called genotypes separately on sex-linked scaffolds by using the “-- haploid” flag. The genotype data was filtered using a combination of vcftools 0.1.15 (Danecek et al. 2011) and vcflib (https://github.com/vcflib/vcflib). First, the raw set of variants was filtered for overlap with annotated repeats. We further filtered variants that had mean site coverage of at least twice the median mean coverage across all sites. For females, this value was calculated separately for autosomal and sex-linked scaffolds. Next, we filtered for sites with a quality score larger than 20 (Q20), for alleles that were supported by at least one read on each strand (SAF>0 & SAR>0) and by at least one read centered to left and right side of the variant (RPL>0 & RPR>0). We further removed male genotypes and female genotypes on autosomes that had a coverage less than 10x. For sex-linked scaffolds in females we set the corresponding coverage threshold to 5x. Following the genotype filtering we removed any sites that had less than 80 % of called genotypes. Finally, we decomposed complex variants and short haplotypes into SNPs and indels and extracted bi-allelic SNPs for downstream analyses. SNPs were intersected with different annotation features using vcfintersect from the vcflib package.

We used vcftools to calculate nucleotide diversity. We downloaded a version of the software from https://github.com/jydu/vcftools, which in contrast to the official release support haploid data for calculations of diversity. For females, the sex-linked scaffolds vcftools were analysed using the “-haploid” flag. Nucleotide diversity was calculated per SNP and summarized across scaffolds or particular scaffold intervals. To get a more unbiased estimate of the nucleotide diversity, we also estimated the number of callable sites in the genome. For this purpose, we used samtools mpileup to output coverage of each sample for each site. The software was run with default settings except for including reads with a minimum mapping quality of 1 and was run separately for males and females. From the raw output we filtered sites based on the same coverage thresholds (depending on sex and autosomal or sex-linked scaffolds) and missingness of genotypes (80 % called genotypes) as employed for the genotype filtering, and further removed positions that were overlapping with annotated repeats. The callable sites were intersected with different annotation features using BEDTools version 2.17.0 (Quinlan & Hall 2010).

## ACCESSIBILITY STATEMENT

All sequence data used for this study will be made accessible under BioProject ID PRJNA578893.

## AUTHOR CONTRIBUTION

Conceived the study: H.S., M.S., H.W. and B.H. Performed analyses: H.S., M.S., E.P.-W., V.E.K, S.P., H.Z., M.L., L.S. and I.B. Wrote the paper with input from all authors: H.S. and B.H.

## ACKNOWLEDGEMENTS

Samples were collected non-destructively and with the appropriate permissions from the Malmö-Lunds djurförsöksetiska nämnd (M 45-14 and 17277-18). Sequencing was performed by the SNP&SEQ Technology Platform in Uppsala, which is part of the NGI Sweden and SciLifeLab, supported by the Swedish Research Council and the Knut and Alice Wallenberg Foundation. Bioinformatics analyses were performed on resources provided by the Swedish National Infrastructure for Computing (SNIC) at Uppsala Multidisciplinary Center for Advanced Computational Science (UPPMAX). V.E.K., E.P.-W. and B.N. are financially supported by the Knut and Alice Wallenberg Foundation as part of the National Bioinformatics Infrastructure Sweden at SciLifeLab. The research was funded by grants from the Royal Physiological Society in Lund, the Erik Philip-Sörensen’s Foundation, Stiftelsen Olle Engkvist and the Wenner-Gren Foundations (to S.P.), the European Research Council under the European Union’s Horizon 2020 research and innovation programme (to H.W., starting grant no. 679799), and the Swedish Research Council (to B.H., consolidator grant no. 621-2016-689).

