## Supplementary method and figures for "Genomics of an avian neo-sex chromosome reveals the evolutionary dynamics of recombination suppression and sex-linked genes"

### Supplementary Methods

#### Configuration file for the falcon assembly:

```
[General]
input_fofn = input.fofn

length_cutoff = 8000
length_cutoff_pr = 12000

target = assembly

job_type = sge
job_queue = falconqueue1
sge_option_da = -pe fpe 4 -q %(job_queue)s
sge_option_la = -pe fpe 4 -q %(job_queue)s
sge_option_cns = -pe fpe 8 -q %(job_queue)s
sge_option_pda = -pe fpe 4 -q %(job_queue)s
sge_option_pla = -pe fpe 4 -q %(job_queue)s
sge_option_fc = -pe fpe 16 -q %(job_queue)s

default_concurrent_jobs = 96

pa_DBsplit_option = -x500 -s400
ovlp_DBsplit_option = -x500 -s400

falcon_sense_option = --output_multi --min_idt 0.70 --min_cov 4 --max_n_read 200 --n_core 8 --min_cov_aln 4 --
min_len_aln 40

overlap_filtering_setting = --max_diff 100 --max_cov 100 --min_cov 1 --n_core 8 --bestn 10

pa_HPCdaligner_option =   -v -B128 -t16 -e.70 -l1000 -s1000 -M28
ovlp_HPCdaligner_option = -v -B128 -t32 -e.96 -l500  -s1000 -M28 -h60

falcon_sense_skip_contained = false
skip_checks = True
dust = false
dazcon = false

use_tmpdir = /scratch
```

### Supplementary Figures

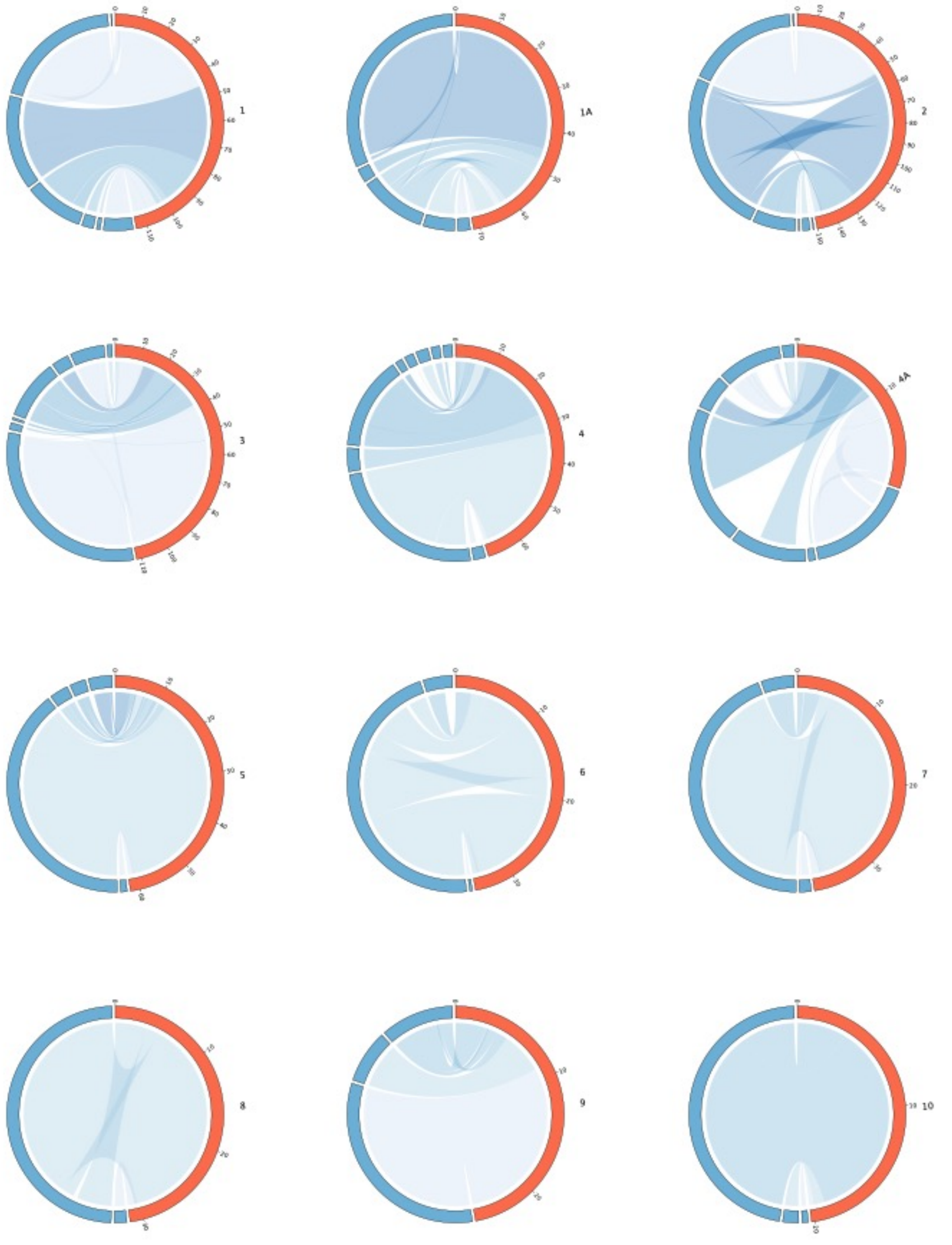


*Supplementary Figure 1a. Synteny plots between great reed warbler scaffolds (blue) and great tit chromosomes (red; with chromosome names).*

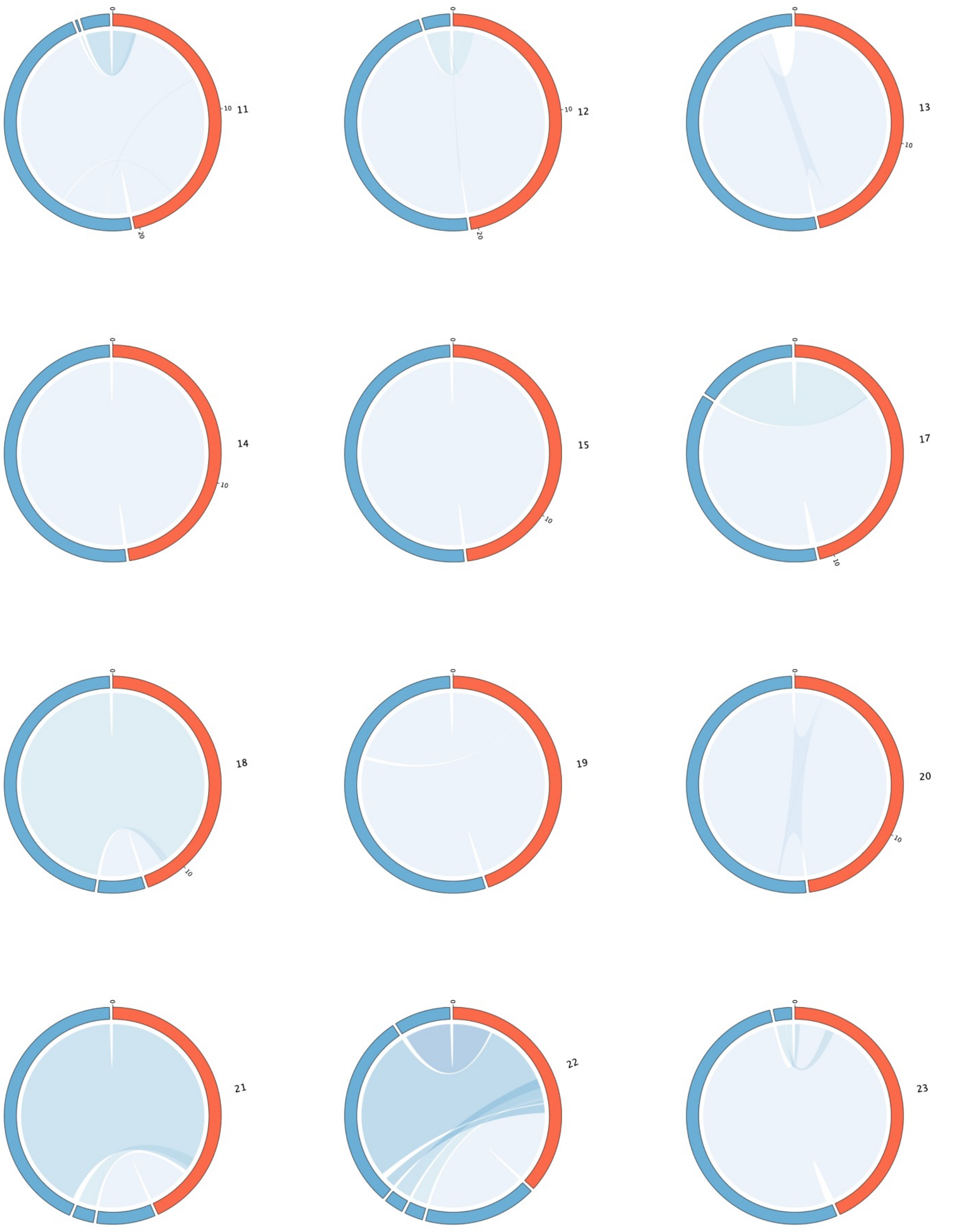


*Supplementary Figure 1b. Synteny plots between great reed warbler scaffolds (blue) and great tit chromosomes (red; with chromosome names).*

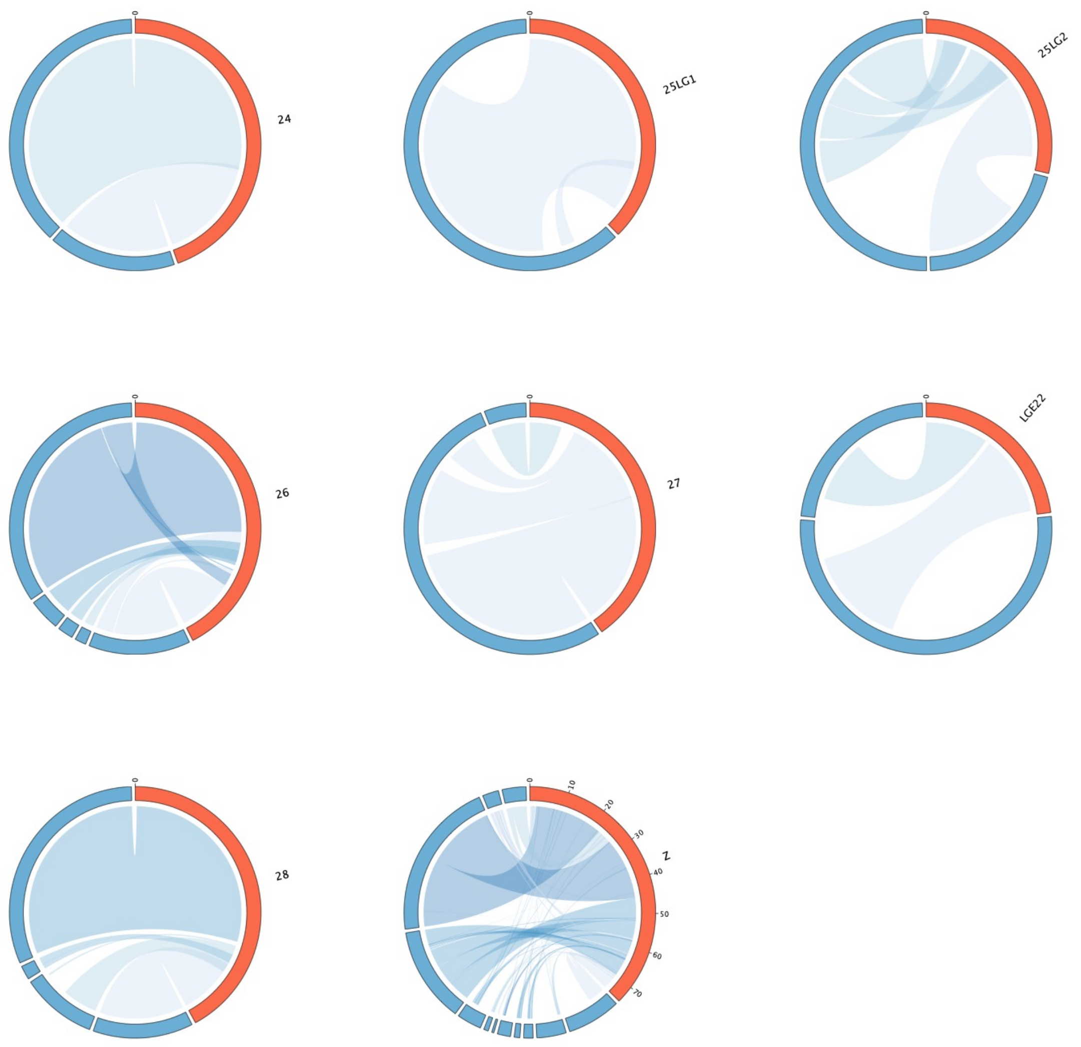


*Supplementary Figure 1c. Synteny plots between great reed warbler scaffolds (blue) and great tit chromosomes (red; with chromosome names).*

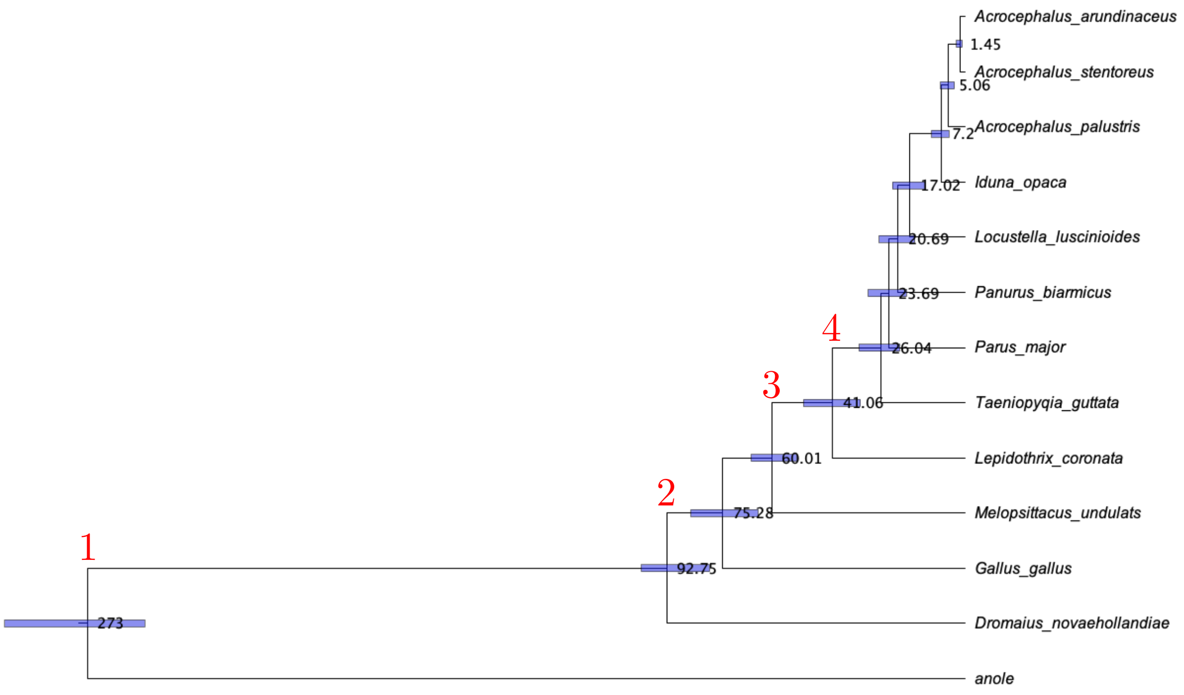


*Supplementary Figure 2. Fossil evidence-dated 13-species phylogeny reconstructed with MCMCTree. Calibrations were denoted with red numbers above nodes, including: (1) 255.9 - 299.8 Myr (Jarvis et al. 2014), (2) 66 - 99.6 Myr. (Jarvis et al. 2014), (3) 51.81 – 66.5 Myr. (Oliveros et al. 2019), (4) 27.25 – 56 Myr. (Oliveros et al. 2019).*

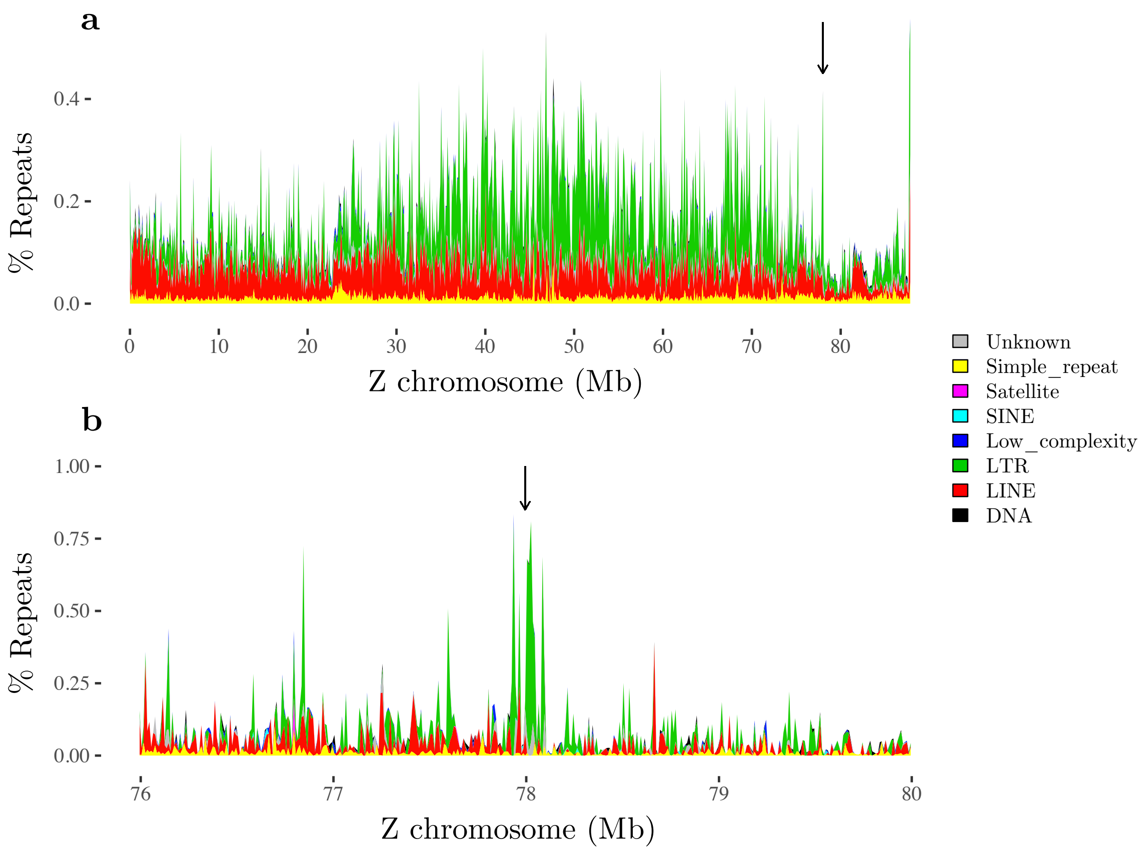


*Supplementary Figure 3. Percentage of repeats along the Z chromosome, colored by type of repeat. The arrow marks the position of the Z to 4A fusion breakpoint. (a) Same as Figure 4b, calculated using window sizes of 100kb. (b) Zoom in on the fusion breakpoint site, calculated using window sizes of 10kb.*

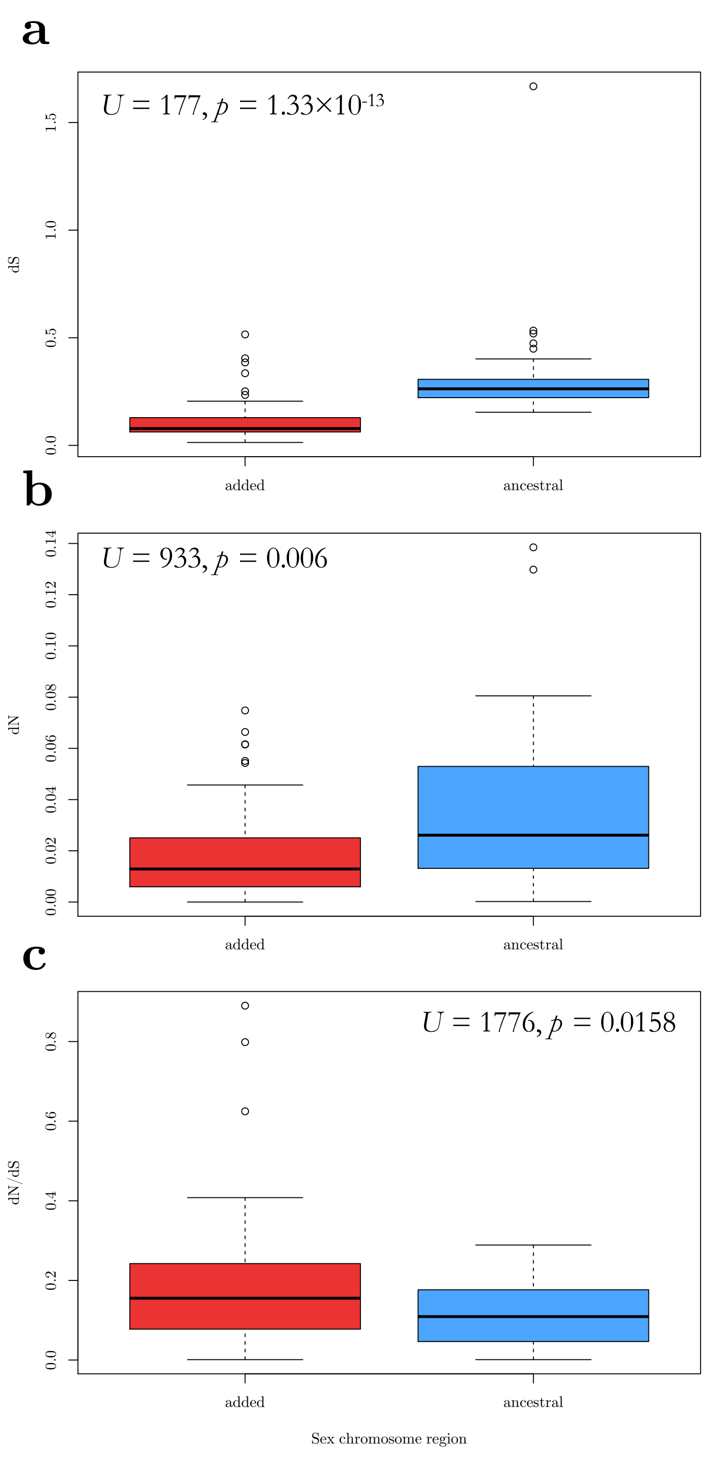


*Supplementary Figure 4. Substitution rates between ZW gametologs positioned on the ancestral (blue) and added (red) sex chromosome region. (a) dS, (b) dN and (c) dN/dS. Statistics from Mann-Whitney U tests are reported for each panel.*

**dN/dS ancestral sex chromosome**

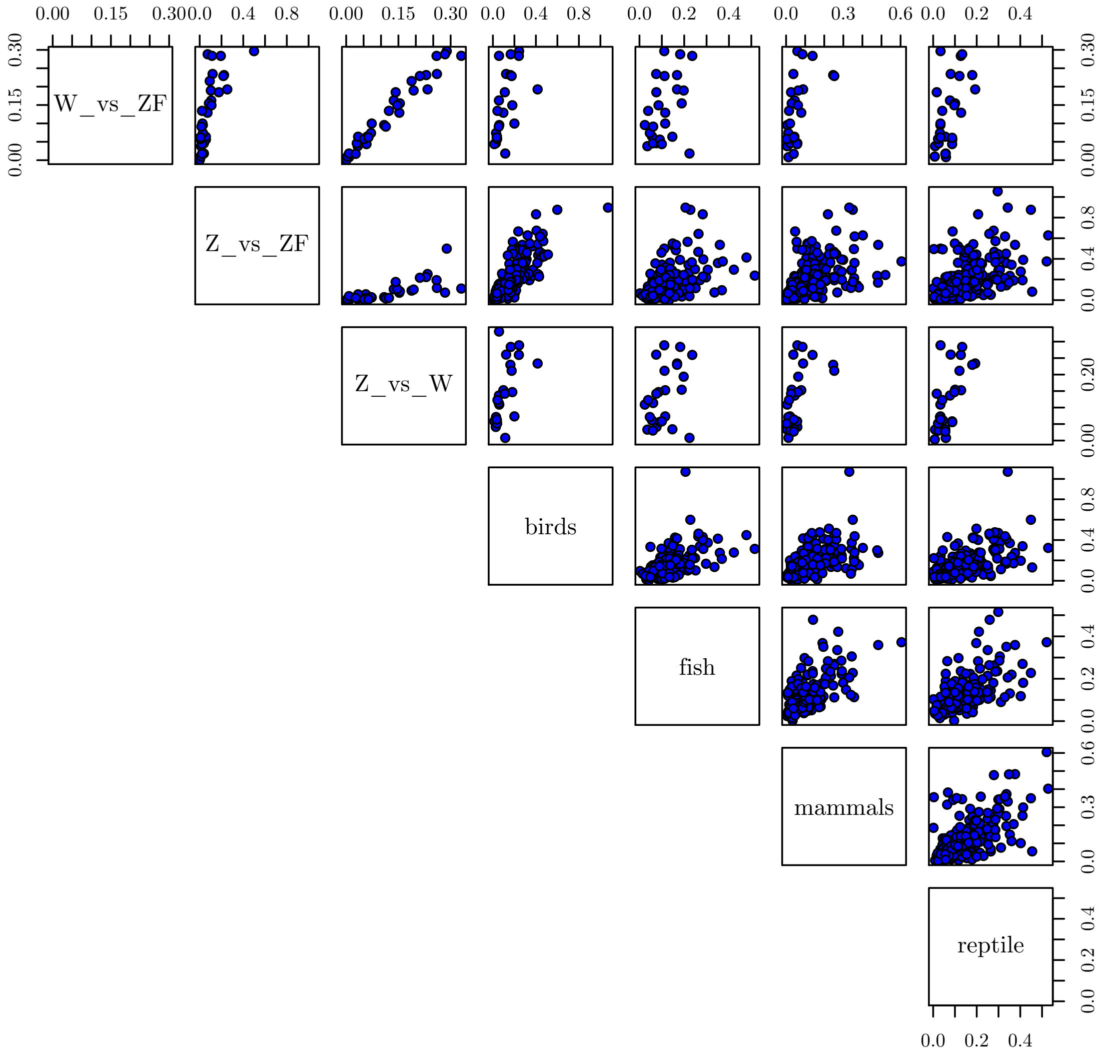


*Supplementary Figure 5. Scatter plots of the dN/dS values used to produce Figure 5a. Comparisons: W_vs_ZF (great reed warbler W vs. zebra finch), Z_vs_ZF (great reed warbler Z vs. zebra finch), birds (chicken vs. zebra finch), fish (stickleback vs. fugu), mammals (human vs. mouse), reptile (green anole vs. bearded dragon).*

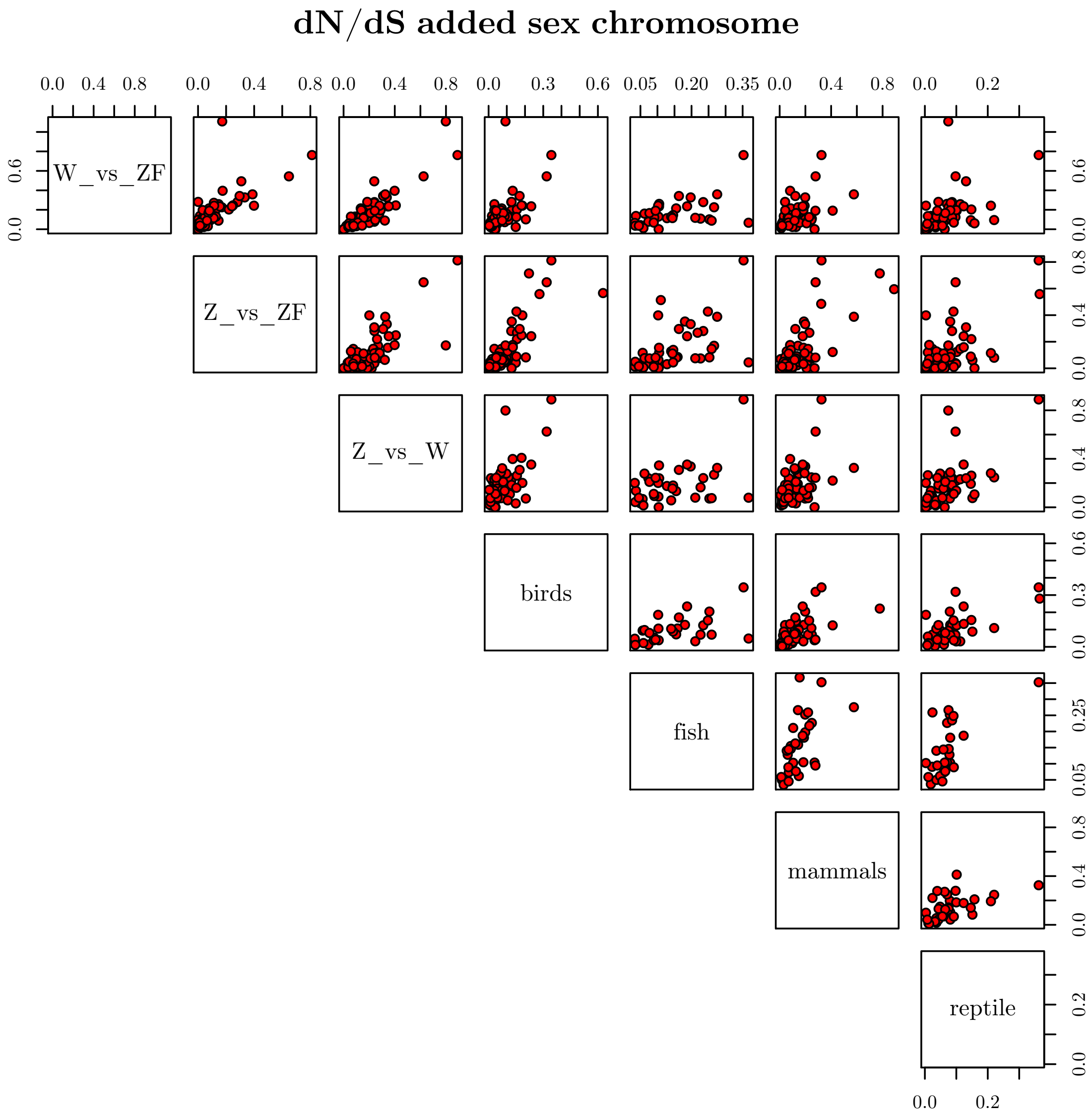


*Supplementary Figure 6. Scatter plots of the dN/dS values used to produce Figure 5b. Comparisons: W_vs_ZF (great reed warbler W vs. zebra finch), Z_vs_ZF (great reed warbler Z vs. zebra finch), birds (chicken vs. zebra finch), fish (stickleback vs. fugu), mammals (human vs. mouse), reptile (green anole vs. bearded dragon).*
